# Scalable and efficient DNA sequencing analysis on different compute infrastructures aiding variant discovery

**DOI:** 10.1101/2023.07.19.549462

**Authors:** Friederike Hanssen, Maxime U. Garcia, Lasse Folkersen, Anders Sune Pedersen, Francesco Lescai, Susanne Jodoin, Edmund Miller, Matthias Seybold, Oskar Wacker, Nicholas Smith, nf-core community, Gisela Gabernet, Sven Nahnsen

## Abstract

DNA variation analysis has become indispensable in many aspects of modern biomedicine, most prominently in the comparison of normal and tumor samples. Thousands of samples are collected in local sequencing efforts and public databases requiring highly scalable, portable, and automated workflows for streamlined processing. Here, we present nf-core/sarek 3, a well-established, comprehensive variant calling and annotation pipeline for germline and somatic samples. It is suitable for any genome with a known reference. We present a full rewrite of the original pipeline showing a significant reduction of storage requirements by using the CRAM format and runtime by increasing intra-sample parallelization. Both are leading to a 70% cost reduction in commercial clouds enabling users to do large-scale and cross-platform data analysis while keeping costs and CO_**2**_ emissions low. The code is available at https://nf-co.re/sarek.

## 1 Introduction

Genomic variation analysis of short-read data has become a key step for modern personalized medicine as well as for fundamental biomedical research. In particular, for biomedical assessment, it is used for characterizing genomes of samples taken from both healthy or tumor tissue. In clinical applications, the resulting information can be used to classify tumors and support treatment decisions [1–3] or research questions, such as drug development [4] or identify variations of interest in larger cohorts for further studies [5, 6]. The technologies and protocols for generating DNA sequencing data vary a lot. Each of the technologies comes with different specialties ranging from targeted gene panels and whole exomes (WES) to whole genomes (WGS) resulting in raw data files from a few to hundreds of gigabytes (GB). Various project-specific factors play a role in choosing the appropriate sequencing technologies, such as the particular type of genomic variations of interest, the cost for sequencing, analysis, and data storage or turn-around times [7]. Panel and exome sequencing is cheaper than WGS [8]. Targeting defined regions allows for having high coverage in these regions. Hence, single nucleotide variants (SNVs) and small insertions and deletions (Indels) can be determined with high confidence. WGS, on the other hand, can be used to additionally investigate more complex alterations such as non-coding variants, large structural variants (SV), and copy-number variations (CNV). Another aspect is ethical considerations on how to handle ‘accidentally detected’ genomic variation in non-targeted genes which had been identified during the whole genome or whole exome sequencing [9, 10].

Examples of large-scale genomics collection projects are TCGA/ICGC or the 100,000 Genomes Project. Some 6,800 whole-genome samples from the former were uniformly processed for the ‘Pan Cancer Analysis of Whole Genomes’ study to obtain a consistent set of somatic mutation calls [11]. More than 12,000 whole-genome samples from the latter were analyzed with respect to their mutational signatures to gain insights into tissue-specific markers [12]. There are several national and international initiatives that aim at gathering more and more sequenced genomes, such as the Estonia Genome Project, the German Human Genome-Phenome Archive, the Iceland Genome Project [13], or the European ‘1+ Million Genomes’ Initiative. Such studies often encompass many patients and their samples are often collected over longer periods of time at multiple sites. This requires stable, and reproducible pipelines that can be run on a variety of different high-performance clusters and cloud setups with differing scheduling system for distributed and homogeneous data processing [14].

Several pipelines [11, 15–18] have been published in different workflow languages to automatically process reads from FastQ files to called (and annotated) variants accompanied by countless in-house workflows. With a certain variety of tools, the workflows usually encompass: quality control steps, read trimming, mapping, duplicate marking, base quality score recalibration, variant calling, and possibly annotation (see Fig. 1).

**Fig. 1:**
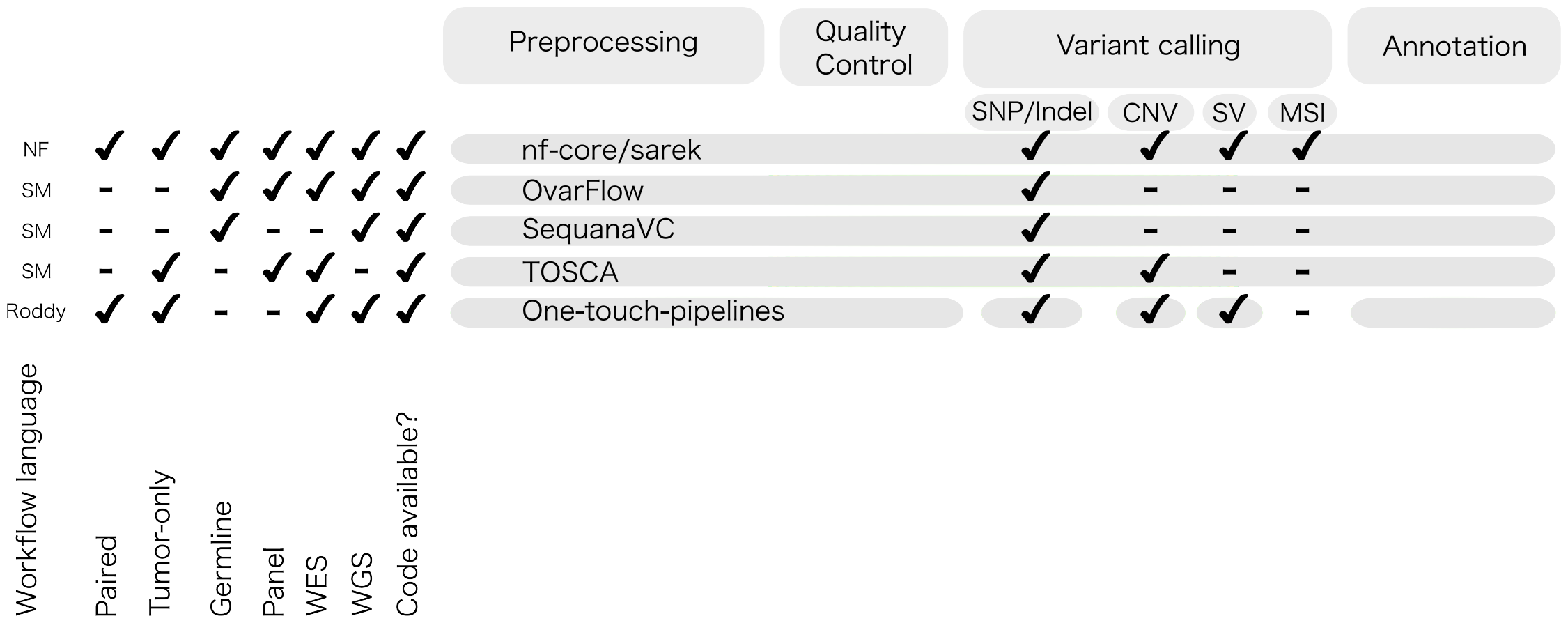
There are many published and countless unpublished variant calling pipelines written in dedicated workflow languages like Nextflow (NF), SnakeMake (SM) [19], and Roddy. All pipelines align the reads, duplicate mark them, and employ various QC metrics (see A1). nf-core/sarek, Ovarflow, and TOSCA have additional option for base quality score recalibration. All pipelines allow variant calling and annotation. The varying supported variant calling types are highlighted for each pipeline respectively. For the One-touch pipelines (OTP), separate workflows have to be triggered for each variant calling type with build-in annotation. nf-core/sarek stands out by covering germline, tumor-only, and paired variant calling followed by annotation across whole genome, whole-exome, and panel sequencing data. The code is available online and well-documented, implemented in Nextflow to enable portability to various infrastructures, and supported by an active community.

While there are many workflows available, nf-core/sarek [15] stands out with its ability to process germline, tumor-only, and paired samples in one run. It can perform SNV/Indel, SV, and CNV calling, as well as micro-satellite instability(MSI) analysis of WGS, WES, and panel data with currently 12 different tools. Since the pipeline is written in Nextflow [20], it benefits from the portability to any supported infrastructure, in particular several cloud vendors and common HPC schedulers enabling cross-platform homogeneous data processing. Furthermore, the pipeline allows the processing of non-model organisms. While the reference genomes and databases are most comprehensively provided for human and mouse genomes as well as subsets for many other organisms, they can be generated and saved for future runs for non-supported species. nf-core/sarek is part of the nf-core community project [21] and has a growing user base with now 242 stars on GitHub and 47.47K unique repository visitors since July 2019 (as of 1st June 2023) who additionally contributes with supporting and improving the code base either by direct contributions, suggesting features, or raising issues.

The pipeline has been used within the field of cancer research [22–27] and beyond, such as the identification of rare variants in tinnitus patients [28], finding SNPs in driver genes related to stress-response in cowpeas [29], the genomic profiling of wild and commercial bumble bee populations [30], or the Personal Genome Project-UK [31]. Here, we present a re-implementation of the nf-core/sarek pipeline using the Nextflow DSL2 framework, an extension of the Nextflow syntax allowing to develop pipelines in a modular fashion, which increases user-based customization to maintain a modern pipeline. The re-implementation is focused on reducing required compute resources for efficient runs on different infrastructures. Minimizing required computing resources has always been of large interest. In particular, in the genomics space, more users run their calculations on several commercial and non-commercial cloud platforms [14]. Commercial platforms usually come with a pay-per-use model, thus there is a high interest to reduce costs due to finite funding. For non-commercial platforms or local clusters, direct costs are possibly of lower interest, however, reducing required resources allows for processing more samples in a shorter time frame. Furthermore, all used tools have been updated to their latest version upon release. For various steps, new tool options have been added, i.e. mapping with DragMap or variant calling with DeepVariant [32], or fastP [33] for adapter trimming.

Using this re-implementation, we show for the first time that population-scale homogeneous recomputing of WGS on commercial clouds is possible.

Our findings demonstrate a 69% reduction in compute costs when utilizing the nf-core/sarek 3.1.1 pipeline, in comparison to a previous version, nf-core/sarek 2.7.2. This translates to costs of just $20 for comprehensive germline short and structural variant calling, and annotation.

## 2 Results

### 2.1 Pipeline overview & summary of new tools and features

An overview of nf-core/sarek v3.1.1 is shown in Fig. 2. The input data is an nf-core community standardized samplesheet in comma-separated value (CSV) format, that provides all relevant metadata needed for the analysis as well as the paths to the FastQ files. The pipeline has multiple entry points to facilitate (re-)computation of specific steps (e.g. recalibration, variant calling, annotation) by providing a samplesheet with paths to the intermediary (recalibrated) BAM/CRAM files. The pipeline processes input sequencing data in FastQ file format based on GATK best-practice recommendations [34], [35]. It consists of four major processing units: pre-processing, variant calling, variant annotation, and quality control (QC) reporting.

**Fig. 2:**
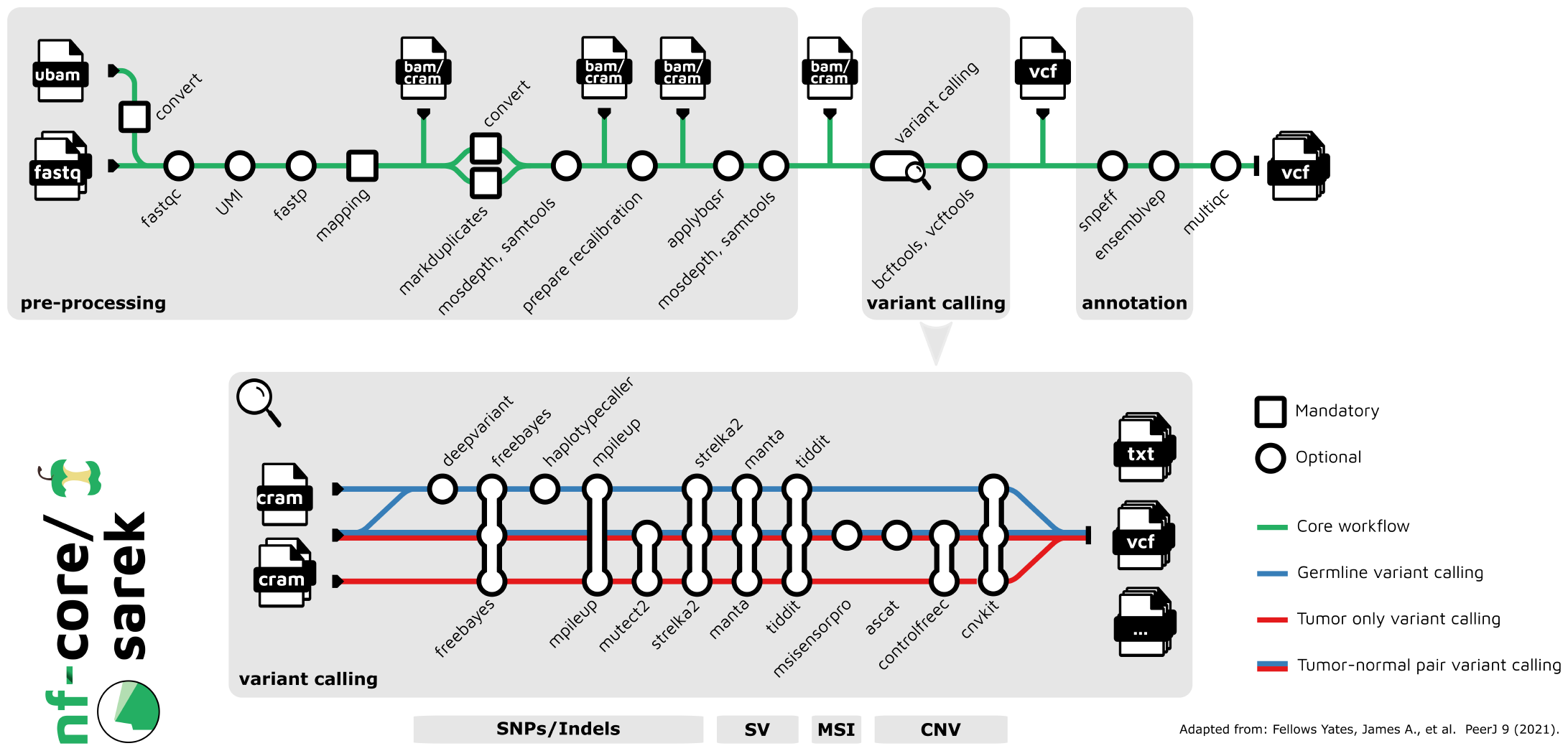
Overview over nf-core/sarek. The pipeline consists of three sections: pre-processing based on GATK Best-practice recommendations (mapping, duplicate marking, and base quality score recalibration), variant calling supporting tools for SNP/Indel, SV, CNV, MSI calling, and annotation. Throughout the pipeline, various quality control tools are run and collated into a comprehensive MultiQC report. The variant calling tools can be mixed in any combination and are all run in parallel.

#### Pre-processing

Enabling homogeneous processing of global genomic resources requires flexibility on the genomic “raw” input data. To cope with the fact that different data repositories provide their “primary” data in different formats, nf-core/sarek support both BAM and FastQ as input. When BAMs are provided as starting input, they are converted to FastQ via SAMtools [36] which allows for fully homogenous processing independent of the provided input format.

The FastQ files are then split into shards with fastP including optional adapter trimming allowing the subsequent alignment step to be run on smaller machines. FastP has been introduced with the release v3.0 and is advantageous over other splitting and adapter removal tools as it combines FastQ sharding and adapter removal into one step, speeding up the computation. With this new implementation, we no longer need to rely on Trim Galore! and Nextflow’s native `splitFastq()` function.

Version 3.1.1 of nf-core/sarek can handle UMI barcodes, which are used in some protocols to detect low allele frequency variants [37]. The user can opt for using Fulcrumgenomics’ fgbio^1^ tool, which generates a consensus read among the ones carrying the same UMI. It will then use these reads as input for the remaining pre-processing steps.

The split FastQ files are aligned with one of the available mappers, which include BWA-MEM [38], BWA-MEM2 [39], or DragMap,^2^ and name-or coordinate-sorted with SAMtools. By adding DragMap support, we comprehensively cover the community’s needs. We added the missing pre-computed reference indices for BWA-MEM2 and DragMap for GRCh38 and GRCh37 to speed up the computation. As recommended by GATK guidelines, we use the entire genome during mapping. Off-target reads for WES and panel analysis are removed according to the provided BED file during the base quality score recalibration (BQSR) step.

By default, the aligned BAMs are then merged, duplicates are marked with GATK4 Markduplicates, and converted to CRAM format in one process to reduce runtime and storage needs. The duplicate marking step was improved by providing name-sorted alignment files to GATK4 MarkDuplicatesSpark. If duplicate marking is skipped, SAMtools is used for merging and conversion to CRAM format. BQSR on the resulting CRAM files is run with GATK4 BaseRecalibrator and GATK4 ApplyBQSR. For both, the GATK Spark implementation is available. Both steps can be skipped, in which case the mapped BAMs are converted to CRAMs using SAMtools.

In order to speed up the computation, genomic regions are processed in parallel following the duplicate marking step. Small regions are grouped and processed together to reduce the number of jobs spun up. By default, interval lists for the complete reference genome provided by GATK are used, enabling scattering by chromosome, and removing unresolved and difficult regions. For targeted sequencing data, we have added support to use the respective target bed files for parallelization as recommended by the GATK guidelines.^3^ Previously, BQSR was always run on the intervals provided for WGS, which led to recalibrating off-target reads increasing computational resources needed. We have added further support to allow users to control group size not just for custom interval files, but also for the ones generated from the genomic regions, allowing a more tailored setup.

#### Variant calling

nf-core/sarek includes a comprehensive set of variant callers to obtain SNPs/Indels, SV, MSI, and/or CNV values using a total of 12 tools (Fig. 2). The variant calling tools have to be selected by the user to ensure the resource footprint is kept low and only necessary tools are run. They are executed in parallel. Newly included tools in the v3.1.1 release are Deepvariant, CNVKit [40], and Tiddit [41]. Furthermore, Haplotypecaller supports both single sample or joint-germline calling [42]. When both Strelka2 [43] and Manta [44] are selected, the candidate Indels from Manta are used for SNP/Indel calling according to the Strelka2 best-practices.^4^ We added a new parameter that allows skipping germline-only variant calling for paired samples to further reduce time, costs, and compute resources for somatic variant calling. Furthermore, scatter-gathering is now supported for all applicable variant calling tools across intervals (see Supplementary Fig. B). The sharded VCF files are then merged with the GATK4 MergeVCF tool. In this way, we reduce computing demands, by avoiding repeated cycles of (de)compressing the files.

#### Variant annotation

The resulting VCF files can be annotated with VEP [45], snpEff [46], or both either separately or by merging the output annotations. The annotation tool VEP has been extended with new plugins allowing more comprehensive annotation: the previously used plugin CADD [47] has been superseded by dbNSFP [48] providing 36 additional prediction algorithms. Furthermore, the plugins LOFTee [49], spliceAI [50], and spliceRegion^5^ have been added.

#### QC reporting

Throughout the pipeline, quality control tools are run, including FastQC before alignment, mosdepth [51], and SAMtools post-duplicate marking and BQSR, as well as vcftools [52] and bcftools on called variants. These results are collected into a MultiQC [53] report together with software version numbers of all the executed tools. The previously used Qualimap [54], which has no direct CRAM support and requires a high amount of computational resources, has been replaced with mosdepth, a fast quality control tool for alignment files. The tool produces comprehensive output files allowing to visualize, e.g. coverage data with the Integrated Genome Viewer (IGV) [55] for easy inspection. In addition, we have enabled quality reporting such as mapping statistics on provided CRAM files, when the pipeline is started from variant calling.

All tools have been updated to their latest stable release at the time of writing. For a complete overview of the most important tool changes, see Supplementary Table C.

#### Pipeline skeleton changes

We expanded the continuous integration testing to include all the new functionality, as well as adding md5sum checking of output files wherever possible. Additionally, we added full size tests that are automatically run on each pipeline release. All nf-core pipelines require a full-size test on realistic data upon release to ensure functionality beyond small test data and portability to cloud infrastructures. The datasets used here include the Genome in a Bottle (GiaB) data set HG001 (down-sampled to 30X WGS) for germline variant calling testing and the tumor-normal pair SRR7890919/SRR7890918 provided by the SEQC2 effort for somatic variant calling testing. Since each of the selected datasets comes with validated VCFs to compare against, they are suited for further benchmarking to investigate the variant calling results. The results for each full size test are displayed on the website nf-co.re/sarek and are available for anyone to explore or download.

High-quality code readability is achieved by combining modules used in the same analysis context into subworkflows, e.g. variant calling with a specific tool and subsequent indexing of the resulting VCF files. In addition, new analysis steps can be added by providing such encapsulated subworkflows, and obsolete parts can quickly be removed entirely. Furthermore, dividing different analysis steps into subworkflows written in separate files simplifies development, which is often done asynchronously with developers at different institutes.

### 2.2 CRAM format allows for storage space reduction

The pipeline has a large data footprint due to the number of computational steps, input data size, and Nextflow’s requirement for a work directory with intermediate results to facilitate resuming. In order to ease storage needs as a possible bottleneck, CRAM files are used as of nf-core/sarek 3. They are a more compressed alternative to BAM files storing only differences to the designated reference. A majority of tools post-duplicate marking support CRAM files. The pipeline can handle both BAM files and CRAM files as in- and output to accommodate various usage scenarios.

We evaluated the resource usage of the two alternatives by running five tumor-normal pairs on nf-core/sarek 3.1.1 as well as on a branch^6^ based on the release in which the internal format was replaced with BAM. Pre-processing (Fig. 3 and Supplementary Fig. D2) and variant calling (Supplementary Fig. D3) were evaluated separately. For eleven processes, the CRAM-based setup resulted in a significant decrease in runtime, for ten an increase in memory, and for eleven an increase in CPU hours could be measured. The overall average CPU hours for the pre-processing benchmark on CRAM version was 3,252.37, in comparison to BAM 3,761.07. The reduction in CPU hours usage, however, had to be compensated by a 34% increase in memory usage. The overall average total memory usage on the CRAM version was 10,346.8GB, for the BAM version it summed up to 7,739.51GB. The storage usage for the work directory for pre-processing these samples drops by 65%, from 170.4TB (BAM) to 59.7TB (CRAM). Processes outputting CRAM files reduce their storage needs by at least a third. In the case of GATK4 ApplyBQSR it was reduced by 64%. Processes operating on CRAM files outputting a different format, e.g. VCFs, show no change in storage usage.

**Fig. 3:**
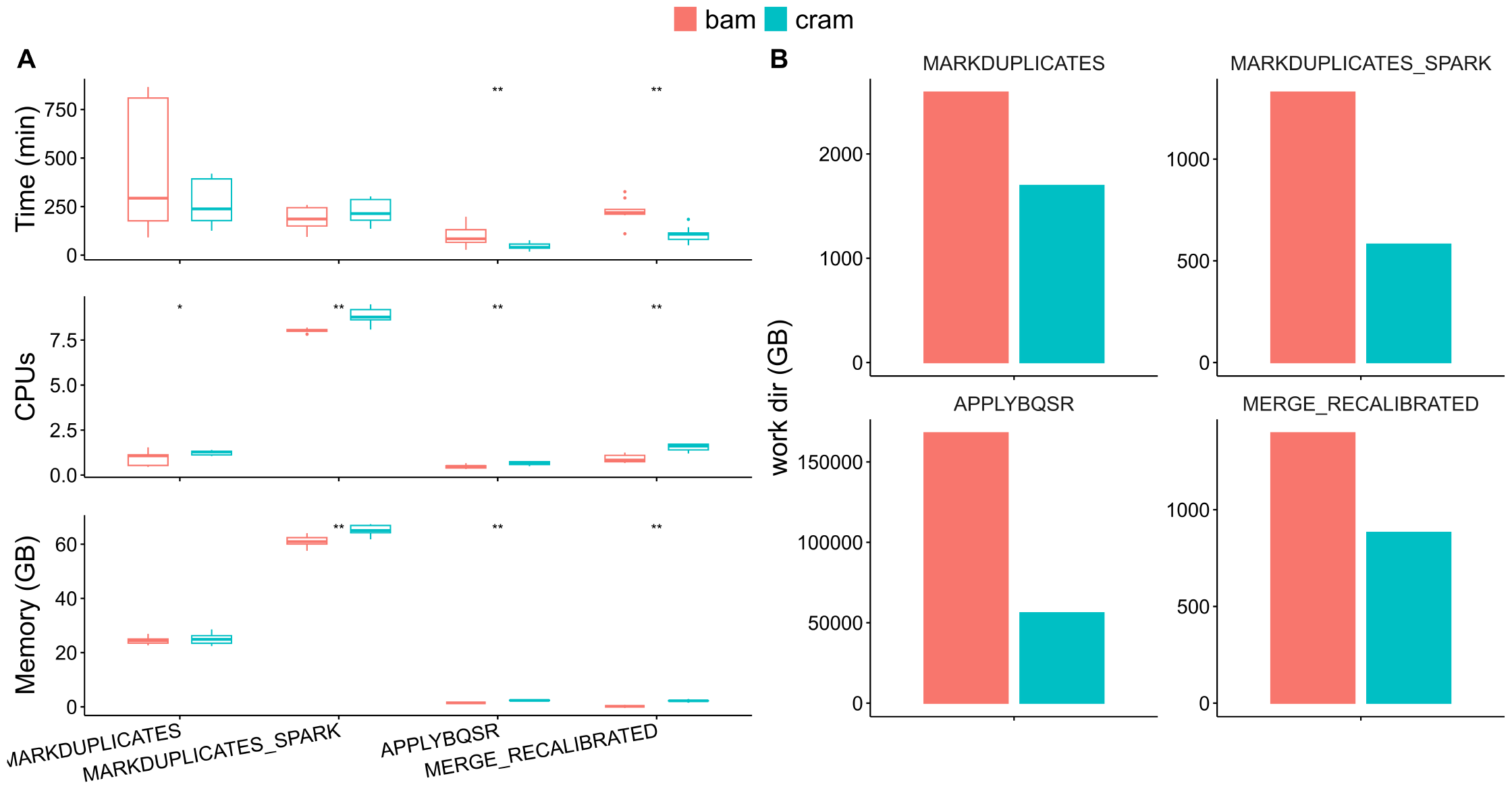
Resource usage of nf-core/sarek 3.1.1 when storing intermediate data in BAM versus CRAM format. **A)** Average realtime, and maximum CPU and memory usage (peak rss) as reported by the Nextflow trace file for the main processes. For processes split within a sample (i.e. ApplyBQSR), the task with the highest runtime per sample is shown as the process runtime. Resource usage was compared using the paired Wilcoxon test (^**^ *p <* 0.01, ^*^ *p <* 0.05). Two out of the four shown processes are significantly faster when using CRAMs instead of BAMs at the expense of an increase in memory or CPU usage. **B)** Storage was evaluated by calculating the total size of the work directories of all tasks of the respective process. Each condition was repeated three times for samples of five tumor-normal paired patients.

### 2.3 Scatter-gather implementations reduce runtime and resource usage

Scatter/gather implementations are highly relevant for parallel processing approaches across genomic regions for BQSR and variant calling. In this release, we have further extended these options: Before mapping, the input FastQ files can now be split and mapped in parallel. For BQSR and variant calling more options to customize the amount of scattering as well as further support for all eligible variant callers are implemented.

We evaluated the impact of different degrees of FastQ file sharding on the mapping process by investigating the division step (fastP), mapping (BWA-MEM), and subsequent merging (GATK4 Markduplicates). The realtime as reported by the Nextflow trace of the longest running mapping process of any one sample was summed up with the realtime of fastP and GATK4 Markduplicates. The space of the work directories of each involved task was summed up, as well as the CPU hours (see Fig. 4A). The overall runtime for the mapping processes decreases until it reaches a plateau at 12 shards, achieving a reduction of the median runtime to 37%. The storage usage increased as soon as any sharding was done due to the sub-FastQs being written to the disk. The CPU hours remain approximately the same due to the long alignment time for large files (see Supplementary Fig E4).

**Fig. 4:**
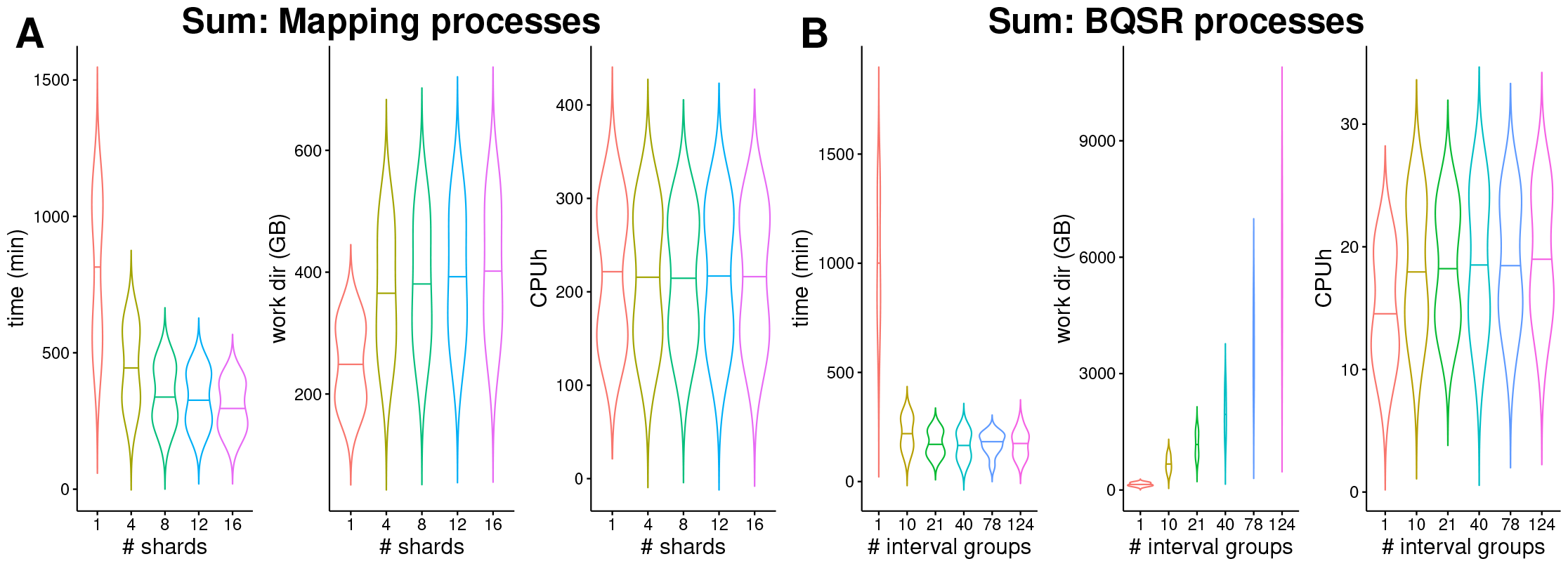
Sharding the input FastQ files and parallelizing computation on interval groups reduces the overall runtime of the nf-core/sarek pipeline. **A)**: Effect of sharding the input files on the mapping processes, including fastP, BWA-MEM, and Markduplicates. The input FastQ files were split into smaller pieces increasing the amounts of shards and the runtime, work directory size, and CPU hours were evaluated for each split size. FastP was run with a different number of CPUs corresponding to the desired number of shards. **B)**: Effect of parallelizing computations across interval groups on BQSR processes, which include the BaseRecalibrator, GatherBQSRReports, Apply-BQSR, and SAMtools merge process. When all intervals were processed together as one group the memory requests for ApplyBQSR had to be increased. The violin plots show computations on tumor-normal paired samples of five patients. The time was evaluated by summing up the highest realtime per task per sample as reported by the Nextflow trace report. The work directory size and CPU hours are the sums of all involved tasks.

We evaluated the impact of different degrees of scattering across genomic intervals on the recalibration and variant calling process with respect to resource usage. Similarly to the mapping processes, the most extended runtime per interval group per samples of all involved processes for BQSR (GATK4 BaseRecalibrator, GATK4 GatherBQSRReports, GATK4 ApplyBQSR, and SAMtools merge) and variant calling (calling and GATK4 MergeVCFs) was summed up respectively. Storage usage and CPU hours results for all tasks were added up. The runtime decreased the most for all measured tools (see Fig. 4B and Supplementary F) when the number of interval groups was set to 21. Raising the number of interval groups did not decrease runtime further. For GATK-based tools, storage usage increased with each further splitting of interval groups. For BQSR storage requirements between 21 intervals groups and 124, the default value, increased by a factor of five. For Deepvariant less storage space was required when applying scattering (Supplementary F6), however, for all other variant callers the storage needs remain on a stable level. The required CPU hours remain stable across various amounts of scattering for all tools.

### 2.4 Benchmarking of short variants against truth VCFs

The pipeline’s SNP and indel variant callers were evaluated for both the germline and paired somatic analysis tracks, the former on three WGS datasets from Genome in a Bottle (HG002-4) [56], the latter on three tumor-normal paired WES datasets from the SEQ2 Consortium [57]. The results were compared to the respective “gold standard” VCFs for high-confidence calls.

We evaluated the precision, recall, and F1 score over all samples. For the germline calls of Deepvariant, only sample HG003 was used since its model was trained on the remaining datasets. The tools’ precision, recall, and F1 scores are in accordance with the previously reported FDA precision challenge [58] results for GiaB samples (see Figure 5A, B for SNPs, Supplementary Figure G8 for Indels). BWA-MEM and BWA-MEM2 lead to higher recall values than DragMap. Strelka2 together with Manta and DeepVariant perform best in all three evaluated metrics. In addition, we investigated one sample sequenced with MGI and BGISEQ respectively with similar results for all variant callers (Supplementary G9, G10).

**Fig. 5:**
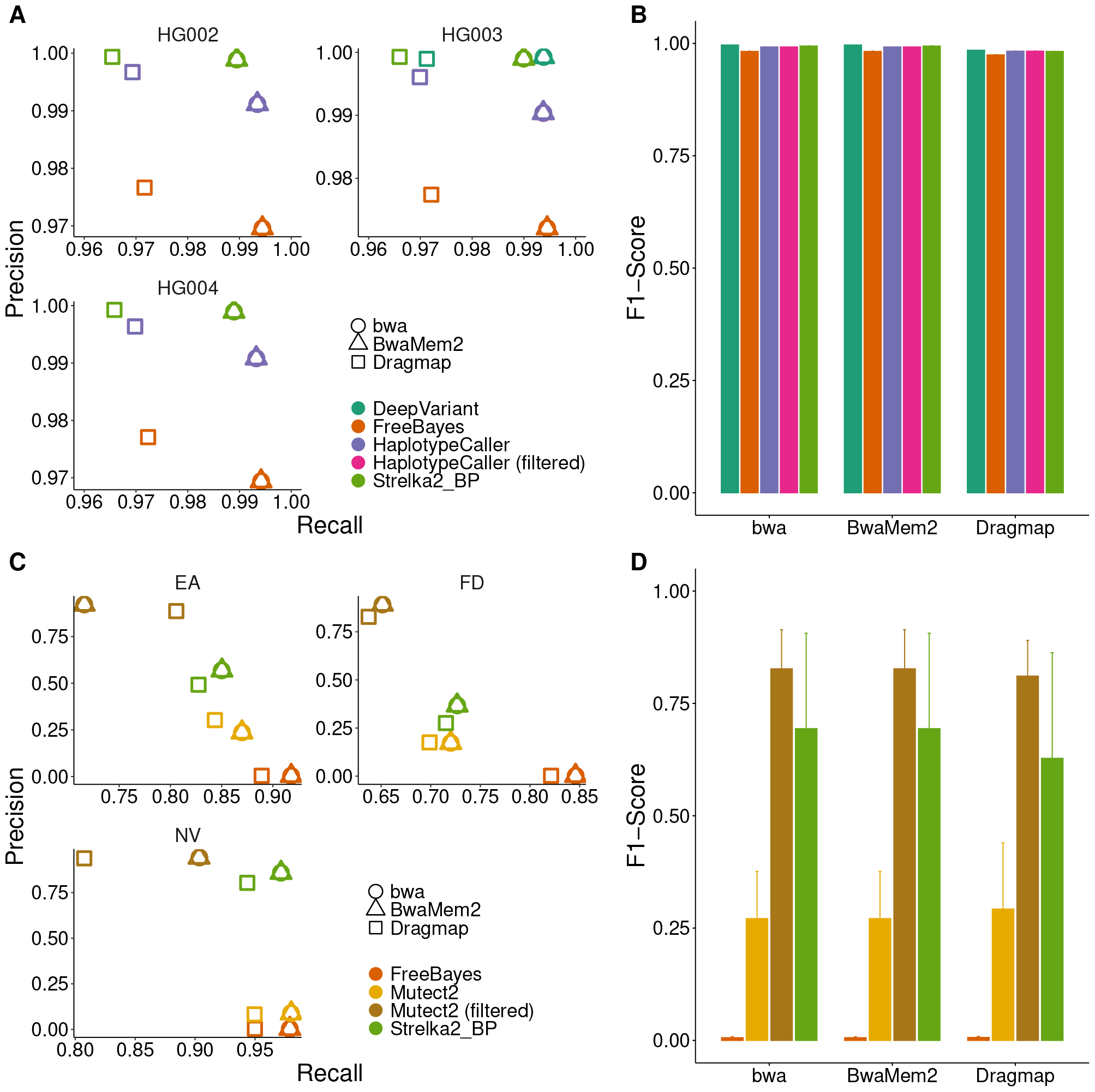
Germline and somatic variant calling evaluation of high-confidence calls using ground-truth benchmarking data with respect to SNPs. **A**,**B**: The germline variant calling track of the pipeline was evaluated using 3 WGS GiaB datasets (HG002-HG004). The average precision, recall, and F1-score values across all the samples are plotted, respectively. **C**,**D**: The somatic paired variant calling track was evaluated using three tumor-normal WES pairs (EA, FD, NV) from SEQ2C.

Similarly, we evaluated the precision, recall, and F1-score for the somatic calls. Filtered Mutect2 calls have the highest precision calls for all samples, and FreeBayes ones the highest recall values (Fig. 5C,D for SNVs). The highest F1-Score is measured for Mutect2, followed by Strelka2. For Indels, Strelka2 outperforms all other tools (Supplementary Figure G8). The results are in-line with what has been previously reported [59].

### 2.5 Comparison of copy-number calls against PCAWG samples

The pipelines’ paired somatic copy number calls from ASCAT, CNVKit, and ControlFREEC were compared against 5 samples from the PCAWG [11] cohort. Each sample has two call sets, one generated with the OTP pipeline and one with the Sanger pipeline (SVCP). In Figure 6, for each tool the calls for each base are evaluated with respect to how many other tools confirmed the base. Calls are divided into two categories: amplifications or deletions. For the former, for patient DO44890 the bases called by each tool are confirmed by at least 3 other tools. Similarly, for DO44919 with an exception for the OTP results, where a set of bases could not be confirmed by any other tool. For a majority of the CNVKit and ControlFREEC calls were confirmed by 3 or more tools for each sample. Overall, there are fewer deletions found than amplifications. For the samples DO44890 and DO44919 a majority of the deletions were called by 2 or more tools. For the remaining samples, calls by CNVKit, ControlFREEC, and SVCP were for a majority of the cases confirmed by one other tool. All calls are visualized in Supplementary H.

**Fig. 6:**
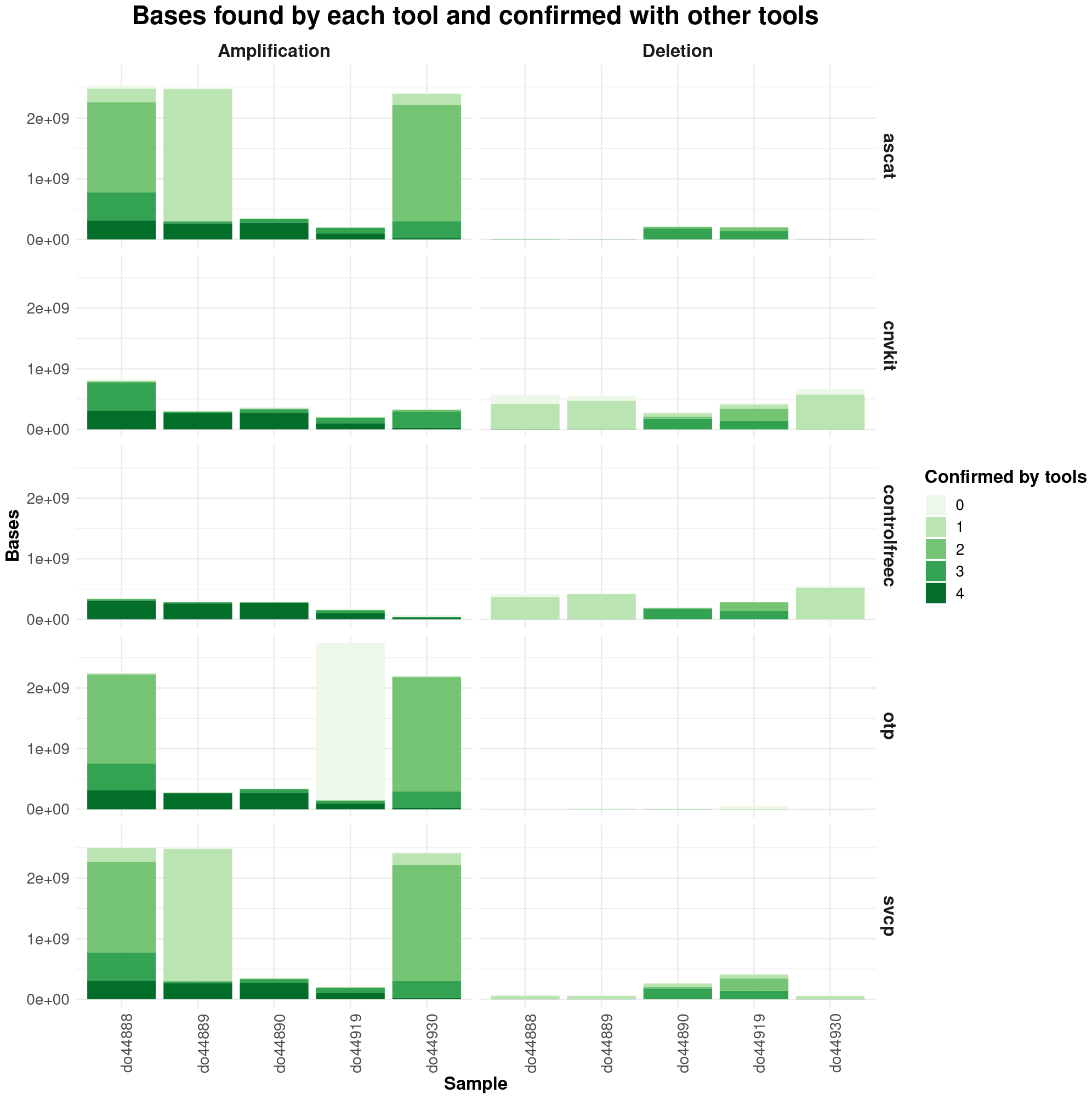
Copy number calling comparison of calls obtained with tools in nf-core/sarek (ASCAT, CNVKit, ControlFREEC) and the two available call sets the ICGC portal for 5 patients from the PCAWG [11] study. The calls are divided into deletions or amplifications. For each event from each caller the number of tools supporting it are plotted.

### 2.6 Portability to AWS and computing costs

The sheer amount of existing genomics data and the unavoidable need for even more data for the detection of disease-causing genotype-phenotype correlations or population-scale analyses will test the limits of on-premise computing sooner or later.

Consequently, more and more data analysis is shifted or supplemented by computation in the cloud. The most recent Nextflow Community survey 2023 indicates that 43% of users use cloud services, which represents an increase of 20% compared to the previous year^7^ with AWS still being the most popular among the respondents. Therefore, we evaluated the cost development of nf-core/sarek between 2.7.1 and 3.1.1 on AWS Batch.

We were able to reduce cloud computing costs to approximately 30%(see Table 1). Furthermore, we could reduce the overall runtime and CPU hours. For a single sample the needed CPU hours are reduced by approximately 70%, and the runtime by 84%. Here, we used spot instances - unused instances auctioned off at a percentage of their on-demand price - whose prices fluctuate constantly. The business models of the cloud providers result in varying spot price percentages and spot prices are frequently subject to change depending on the overall demand.

**Table 1:** Average costs per patient on AWS batch for nf-core/sarek version 2.7.2 and 3.1.1. All pipeline runs were performed with the tools, Strelka2, Manta, and VEP. Each analysis run is repeated three times and run on data from the donor DO50970 with either their normal or tumor-normal paired sample. The normal sample has a median coverage of 36X, the tumor sample of 65X.

| Version | Samples | Avg. costs [\$] | Runtime | CPU hours |
| --- | --- | --- | --- | --- |
| 2.7.2 | 1 normal | 68.04 | 46h8m | 1118.4 |
| 3.1.1 | 1 normal | 20.82 | 12h4m | 342.5 |
| 3.1.1 | 1 paired | 66.83 | 31h47m | 1324.3 |

## 3 Discussion

An essential aspect of high-throughput processing is seamless scalability, one of the main advantages of using a dedicated workflow language such as Nextflow. Combining adapter trimming, quality control, and sharding of FastQ files in a single step, as well as more tailored splitting into intervals for variant calling, reduces the needed CPU hours by 66%. Replacing Nextflow’s native `splitFastq()` function with a dedicated process, allows us to make use of all advantages of regular job submission, including assigning resources to the jobs, automatic retries on job failure, and resume functionality. Previous pipeline versions have typically not been able to split the FastQ files and have, thus, missed out on scalability options. Our experiments have shown clear limits for parallelization into interval groups. There is no further benefit to reducing runtime beyond 21 interval groups. However, storage usage increases. This is respected in future releases by setting the default number of interval groups to 21 further reducing the storage footprint of the pipeline.

The switch to using CRAM files results in reduced storage space usage of 65%, at the expense of higher memory requirements. The additional memory needs are distributed over the thousands of tasks run for the benchmark. Due to this, in practice, memory is not a limiting factor. The additional needs are only required for the task’s run time, usually comprised of a couple of GB at most, and can subsequently be reused. The storage, however, accumulates over the entire pipeline run and can therefore pose a bottleneck for usage scenarios with large input data sets relative to the available storage on the respective compute system.

Both changes, switching to CRAM format reducing storage requirements by two-thirds and reducing the amount of scattering - further reducing storage requirements by a factor of 5, will enable users to run the pipeline on smaller systems more efficiently. While there are many benefits for the scientists running the pipeline, we would like to emphasize also the ecological need to reduce the carbon footprint in computational research. This is relevant in particular considering the ever-growing number of available samples in national and international genome repositories, which aim at facilitating truly comprehensive population-based analysis and understanding underlying genome variations. Our results provide solutions to reduce CO_2_ emissions.

One of the main objectives of this work is to enable scientists to run the pipeline in cloud environments at low costs per sample. Cost-efficient cloud computing is increasingly important for data-driven science. Intransparent and unpredictable cost models discourage a scientist. Using the cloud setup as described in this work, we reduced the costs to 50% in comparison to previous releases. Recently, Seqera Labs posted a blog article^8^ showing further cost reductions when using nonvolatile memory express (NVME) storage, a protocol to accelerate transfer speed, and a new fusion file system promising even cheaper runs in the future. In addition, users can further reduce their cloud costs by selecting compute instances in cheaper AWS regions and fixing the spot price percentage they are willing to pay more tightly by enforcing an upper bound for costs per sample at the expense of possible waiting time for such machines to become available. nf-core/sarek is an established, comprehensive variant calling pipeline in the genomics field, which can be applied to any organism for which a reference genome exists. Future releases further simplify analyses with custom references by enabling pre-computation of all needed indices, an interesting feature when multiple users work with organisms on shared systems for which reference files are not provided by default. On request, such references and their corresponding indices and database files can be added to the central resource AWS-iGenomes and made available to the community.

We benchmarked the pipelines performance for germline and somatic small variants against given truth datasets, as well as comparing copy number calls to ones obtained by the PCAWG study. The copy number evaluation highlights the need for validating calls with multiple tools. There is a set of bases showing strong evidence by being called by all tools. However, some tools, CNVKit and ControlFREEC, show a seemingly more conservative approach with calls validated by almost all others, whereas ASCAT, SVCP, and OTP generated overall more calls which were confirmed by fewer tools. The tools’ performance differs between samples indicating possible further factors need to be taken into account. The similarities between ASCAT and SVCP can be explained by the fact that the SVCP pipeline also uses an earlier version of the ASCAT tool.

The nf-core/sarek rewrite to DSL2 makes the code base more maintainable and easier to read, a factor that is crucial to allow new developers to join the effort with a reasonable learning curve. All pipeline processes are specified in separate files in the form of modules a majority of which are maintained by the nf-core community. Tools used in the same context are combined into subworkflows. They will be added to the nf-core subworkflows collection in the near future allowing further collaboration and shared maintenance across pipelines and beyond the nf-core community. Modularising all tools will enable us to simply do a drop-in replacement when tools should be exchanged for a different one or new ones added as they emerge. nf-core/tools installs them in the appropriate directory, they just need to be called at the appropriate position in a (sub)workflow. Furthermore, the use of modules allows users to customize the released pipeline version at runtime. Before this change, it was necessary to change the underlying code if arguments of a tool were not exposed to the pipeline. This was limiting for users since they had to wait for the feature to be implemented and released before using it. With the new modular config files, arguments can be modified by providing a user-based custom configuration setting the exact command line arguments without changing the underlying code or the release tag. This allows a higher degree of flexibility for the analysis whilst simultaneously using a released version. Reproducibility can then be ensured as before by providing the exact release version, pipeline parameters, and the respective custom config(s).

## 4 Methods

### 4.1 Implementation

nf-core/sarek is a Nextflow-based pipeline that has been part of the nf-core project since release 2.5. Thus, nf-core/sarek is based on the nf-core template, which provides a code and documentation skeleton to ensure current best practices. The pipeline was one of the first to be ported from Nextflow’s domain specific language version 1 (DSL1) to DSL2. The DSL2 framework allows modularization and code sharing. 78 of 80 modules used in nf-core/sarek have been made available in the nf-core community’s shared repository, nf-core/modules, implementing Nextflow wrappers around ideally individual tools. The tools are typically accessible through (bio)conda [60] and have a corresponding docker and singularity container provided by the Biocontainers [61] community enabling portability and reproducibility for each such ‘module’. This single-tool-per-process approach ensures that previously occurring dependency conflicts are mitigated. nf-core/tools, a helper tool for users and developers, allows easy creation, installation, and re-use of these modules, which will be important for further extensions of nf-core/sarek.

### 4.2 Data sets & compute environments

In order to evaluate the computational requirements of the pipeline, five tumor-normal paired samples from the ICGC LICA-FR [62] cohort are used (see Table 2). The unaligned BAM files are downloaded and converted to paired-end FastQ files using nf-core/bamtofastq v1.0.0 (formerly qbic-pipelines/bamtofastq) with Nextflow version 20.10.0 and singularity.

**Table 2:**
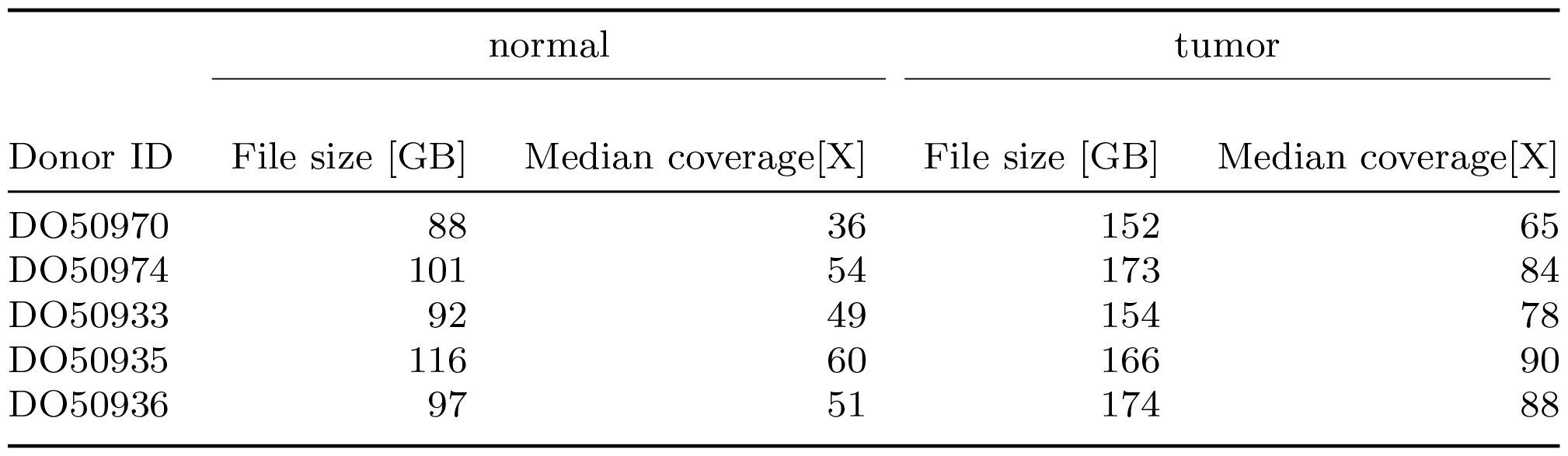
Datasets used for benchmarking are part of the LICA-FR cohort. The BAM files are downloaded and converted to FastQ files. The respective donor IDs and file sizes of the converted FastQ files are listed below.

| Donor ID | normal |  | tumor |  |
| --- | --- | --- | --- | --- |
|  | File size [GB] | Median coverage[X] | File size [GB] | Median coverage[X] |
| DO50970 | 88 | 36 | 152 | 65 |
| DO50974 | 101 | 54 | 173 | 84 |
| DO50933 | 92 | 49 | 154 | 78 |
| DO50935 | 116 | 60 | 166 | 90 |
| DO50936 | 97 | 51 | 174 | 88 |

Unless otherwise indicated, evaluations are done with Nextflow version 22.10.2 build 5832 and Singularity 3.8.7-1.e18 on a shared HPC cluster. A parallel BeeGFS filesystem [63] is used with 1 metadata and 2 storage nodes. Each storage node has 2 raid systems with 10^*^14TB disks respectively. The data systems are connected with a 50GB ethernet connection. The HPC is using Slurm as scheduler and consists of 24 nodes with 32 cores and 64 threads each (2^*^ AMD EPYC 7343) with 512GB RAM and 2TB NVMEe disks as well as four nodes with 64 cores and 128 threads each (2^*^ AMD EPYC 7513) with 20248GB and RAM and 4TB NVMe disks, these NVMes are utilized via Nextflow scratch option. To increase the speed and decrease the load on the filesystem, calculations are therefore performed directly on the NVMe, and only the results are written back to the BeeGFS. The cluster is shared and resources per user are allocated by a Fair-share policy. At any time 100 tasks can be run at most by a user in parallel.

All jobs are submitted using `-profile cfc` providing a cluster-specific configuration, which is stored in the GitHub repository nf-core/configs.^9^

Resource usage for all experiments was evaluated by supplying:

Listing 1: trace.config

~~~
trace {
   fields = ‘task_id, hash, native_id, process, tag, name,
   status, exit, module, container, cpus, time, disk, memory,
   attempt, submit, start, complete, duration, realtime,
   queue,% cpu,% mem, rss, vmem, peak_rss, peak_vmem,
   rchar, wchar, syscr, syscw, read_bytes, write_bytes,
   vol_ctxt, inv_ctxt, workdir, scratch, error_action ‘
   raw = true
}
~~~

in a custom configuration file and collecting the file sizes of the work directory with:

Listing 2: storage.sh

~~~
du - hb -- all -- max - depth =4 < absolute / path / to >/ work /
    > folder_sizes. tsv
~~~

### 4.3 Reducing storage requirements

In this release, the internal file format following duplicate marking is changed to using CRAM files. To evaluate the required compute resources and the actual data footprint for both file formats, five paired ICGC genomes are run through all tools part of nf-core/sarek 3.1.1 and an altered version of nf-core/sarek 3.1.1 that uses the BAM format instead. For each process, singularity containers from the Biocontainers registry are used. Each run configuration is repeated three times. Unless otherwise specified, the default parameters for nf-core/sarek 3.1.1 are used. We evaluate the pre-processing and variant calling steps independently.

We evaluate all processes corresponding to FastQ quality control, aligning the reads to the reference genome, duplicate marking, base quality score recalibration (BQSR), and quality control of aligned reads. The command for running the pre-processing steps is the following:

Listing 3: Pre-processing

~~~
nextflow run nf - core / sarek -r 3.1.1 - profile cfc \
-- input ./ input. csv
~~~

The memory requirements for BWA-MEM are increased to 60GB, as well as the runtime for GATK4 Markduplicates and SAMtools merge to 16h and 8h respectively from the provided defaults.

Secondly, we evaluate variant calling with all tools for all germline and paired somatic variant calling for all samples:

Listing 4: Variant calling

~~~
nextflow run nf - core / sarek -r 3.1.1 - profile cfc \
-- input recalibrated . csv \
-- tools deepvariant, haplotypecaller, mutect2, strelka, \
freebayes, ascat, controlfreec, \
cnvkit, manta, tiddit, msisensorpro \
-- step variant_calling
~~~

The requested time for all processes is increased to 144h to mitigate interruptions due to runtime time-outs by providing a custom configuration file.

### 4.4 Evaluation of pipeline runtime and resource usage

In order to evaluate the impact on runtime and storage requirements, ten samples are run with different sizes of scattered groups: 1, 10, 21, 40, 78, and 124 (default). In order to generate the respective interval group sizes the parameter `--nucleotides_per_second` is set to 5000000, 400000, 200000, 70000, 10001, 1000 (default). For read splitting, fastP is run with various numbers of CPUs specified (respectively, 0, 4, 8, 12 (default), and 16) since the tools generates chunks firstly by number of CPUs and secondly by the maximum number of entries per file as defined by a parameter. Here, it is set to 500,000,000 to prevent any further subdivision.

Example command for 40 interval groups and 8 FastQ file chunks:

Listing 5: Intra-sample parallelization

~~~
nextflow run nf - core / sarek -r 3.1.1 - profile cfc \
-- input input. csv \
-- tools deepvariant, haplotypecaller, mutect2, strelka,
freebayes, ascat, controlfreec,
cnvkit, manta, tiddit, msisensorpro \
-- nucleotides_per_second 70000 \
-- split_fastq 500000000
-c ressource . config
~~~

The memory requirements for BWA-MEM are increased to 60GB for all tests. The memory for FreeBayes is increased to 24GB, for one interval group it is reduced again to 18GB. For GATK4 ApplyBQSR the memory is increased to 16GB when all intervals were processed in one group. For a full list of configs, see https://github.com/qbic-projects/QSARK.

### 4.5 Benchmarking of short variants against truth VCFs

In order to benchmark the variants called by the pipeline, both the germline and paired variant calling tools are evaluated on three datasets each: for the germline callers the GiaB datasets HG002-HG004 [56] are used, for the paired somatic callers three WES datasets from the Sequencing Quality Control Phase II Consortium [57]. Comparisons are made only in the high-confidence regions defined for the benchmark, i.e. omitting difficult-to-call regions.

#### Germline variant calling

The whole-genome germline GiaB samples from an Illumina NovaSeq are downloaded from the GiaB consortium’s ftp-server. Downsampling to 40x is done using the seqtk^10^ tool. nf-core/sarek is run with default parameters. The parameter `--nucleotides_per_second` is increased to 200,000. All eligible variant callers are combined with all three mappers. Comparisons are calculated using hap.py,^11^ version v0.3.14:

Listing 6: Evaluation of germline calls

~~~
hap . py HG00 {2,3,4} _GRCh 38 _1 _22 _v 4 .2.1 _benchmark . vcf. gz \
       query . vcf. gz \
       -o results/ \
       -V -engine = vcfeval \
       -- engine - vcfeval - template grch 38 . sdf \
       -- threads 3 \
       -f \
       HG00 {2,3,4} _GRCh 38 _1 _22 _v 4 .2.1 _benchmark_noinconsistent . bed \
       -- logfile results/
       -- scratch - prefix .
~~~

Short variant calls from Haplotypecaller, Deepvariant, Freebayes, and Strelka2 mapped with Dragmap (base quality recalibration is skipped), BWA-MEM, and BWA-MEM2 are included in the analysis. Deepvariant is evaluated only on HG003.

Furthermore, reads for the sample HG002 sequenced with MGISEQ and BGISEQ500 are downloaded from the manufacturer. They are downsampled to 20X, 30X, 40X, and 50X using seqtk and subsequently processed with nf-core/sarek 3.1.1 using default parameters and all eligible variant callers. Evaluation against the truth VCF is done using hap.py.

#### Somatic variant calling

The somatic short variant calls are evaluated on three whole-exome sequencing datasets: SRR7890919/SRR7890918 (EA), SRR7890878/SRR7890877 (FD), SRR7890830/SRR7890846 (NV) run on an Illumina HiSeq 1500 (EA), 4000 (FD), 2500 (NV). The data is downloaded using nf-core/fetchngs v1.10.0. nf-core/sarek is run on default parameters. In addition, trimming is enabled with `--trim_fastq`. When using DragMap `--skip_tools baserecalibrator` is set. The VCFs are PASS filtered using bcftools v1.10.2. The calls are evaluated using RTGTools [64] for a combination of each mapper with all available variant callers: Strelka2 together with Manta, Freebayes, and Mutect2.

Listing 7: Evaluation of somatic calls

~~~
bcftools view -f ‘PASS,.’ \
        results. vcf. gz \
        -o query . vcf
rtg vcfeval -c query . vcf. gz \
    -b high - confidence_sSNV_in_HC_regions_v 1 .2. vcf. gz \
    -o ./ out/ \
    -t grch 38 . sdf \
    -e High - Confidence_Regions_v 1 .2. bed \
    -- squash - ploidy -- all - records -- sample = ALT \
    -- bed - regions \
    S07604624 _Padded_Agilent_Sure Select XT_allexons_V 6 _UTR . bed
~~~

### 4.6 Comparison of copy number calls against PCAWG samples

In order to evaluate the copy number calls, WGS alignment files for 5 PCAWG patients (DO44888, DO44930, DO44890, DO44919, DO44889) were downloaded and reprocessed with nf-core/sarek. In addition, the provided copy number calls were downloaded from the ICGC portal for each patient. The calls are compared by dividing the calls from each tool into two groups: amplifications and deletions. For each base, it is then determined how many other tools identified for a given the same group. Visualization of the respectively called copy numbers is done using `karyoploteR` [65] and `CopyNumberPlot` [66].

Listing 8: Parameters to run nf-core/sarek for evaluation of copy number calls

~~~
nextflow run nf - core / sarek -r 3.4.0 - profile cfc \
 -- input input. csv -- outdir results \
 -- tools ascat, controlfreec, cnvkit \
 -- only_paired_variant_calling \
 -c ressources_cnv . config
~~~

### 4.7 Portability to AWS cloud and computing cost

In order to evaluate the costs for running nf-core/sarek on AWS batch, the compute environments are created using Tower Forge,^12^ with the following settings: spot instance, max CPUs 1,000, EBS Auto scale, and fusion mounts enabled. Instance types are chosen by using strategy ‘optimal’ with the allocation strategy `spot_capacity_optimized`. The computation is run in AWS region us-east-1. The pipeline runs are launched with Nextflow Tower setting the Nextflow version to 22.10.3 by adding export `NXF_VER=22`.10.3 to the pre-run script and `process.afterScript = ‘sleep 60’` to the config section.

The pipeline is run on the normal sample for DO50970 using default settings together with Strelka2, Manta, and VEP. For one paired sample evaluation, the pipeline was run the tumor-normal pair (DO50970/DO50970). nf-core/sarek 3.1.1 is run with `--nucleotides_per_second` 200000. For the paired run, we set `--only_paired_variant_calling`.

The costs for nf-core/sarek 2.7.1 are evaluated on the same normal sample using Strelka2, Manta, and VEP on default parameters. Compute resources for mapping (372GB memory, 48 cpus), duplicate marking (30GB memory, 6 cpus), quality control with BamQC (372GB memory, 48 cpus), GATK4 BaseRecalibrator (4GB memory, 4 cpus), and GATK4 ApplyBQSR (4GB memory) is increased.

## Supplementary information

All supplementary files are available at https://github.com/qbic-projects/qsark.

## Acknowledgments

We want to thank Johannes Köster for his advice during the benchmarking. We would also like to thank the nf-core and Nextflow community for developing the nf-core infrastructure and resources for Nextflow pipelines and, in particular Mahesh Binzer-Panchal and Alexander Peltzer, for their assistance in reviewing and releasing the pipeline. A full list of nf-core community members is available at https://nf-co.re/community.

## Declarations

### Ethics approval

Not applicable.

### Consent to participate

Not applicable.

### Consent for publication

All authors read the manuscript and consent to its publication.

### Availability of data and materials

The comparison pipeline using BAM files is available at https://github.com/FriederikeHanssen/sarek/tree/bam31. All pipeline run commands used for the benchmarking, configurations, trace reports, evaluation, and visualization scripts are available at https://github.com/qbic-projects/qsark.

### Code availability

The pipeline is available at https://github.com/nf-core/sarek and each release is archived on Zenodo https://doi.org/10.5281/zenodo.3476425

### Authors’ contributions

FH and MUG lead the project. FH, MUG, LF, ASP, FL, SJ, OW, NS wrote the pipeline. GG added the full-size tests. EM and MUG added the CI testing framework. LF and FH added the evaluation against the truth datasets. GG and FH added the AWS experiments. FH added the remaining experiments. MS maintains the core facility cluster and supported the runtime benchmarking experiments. FH, LF, and FL wrote the manuscript with comments from all authors. GG and SN supervised the project.

## Appendix A Overview of quality control tools in variant calling pipelines

**Table A1:**
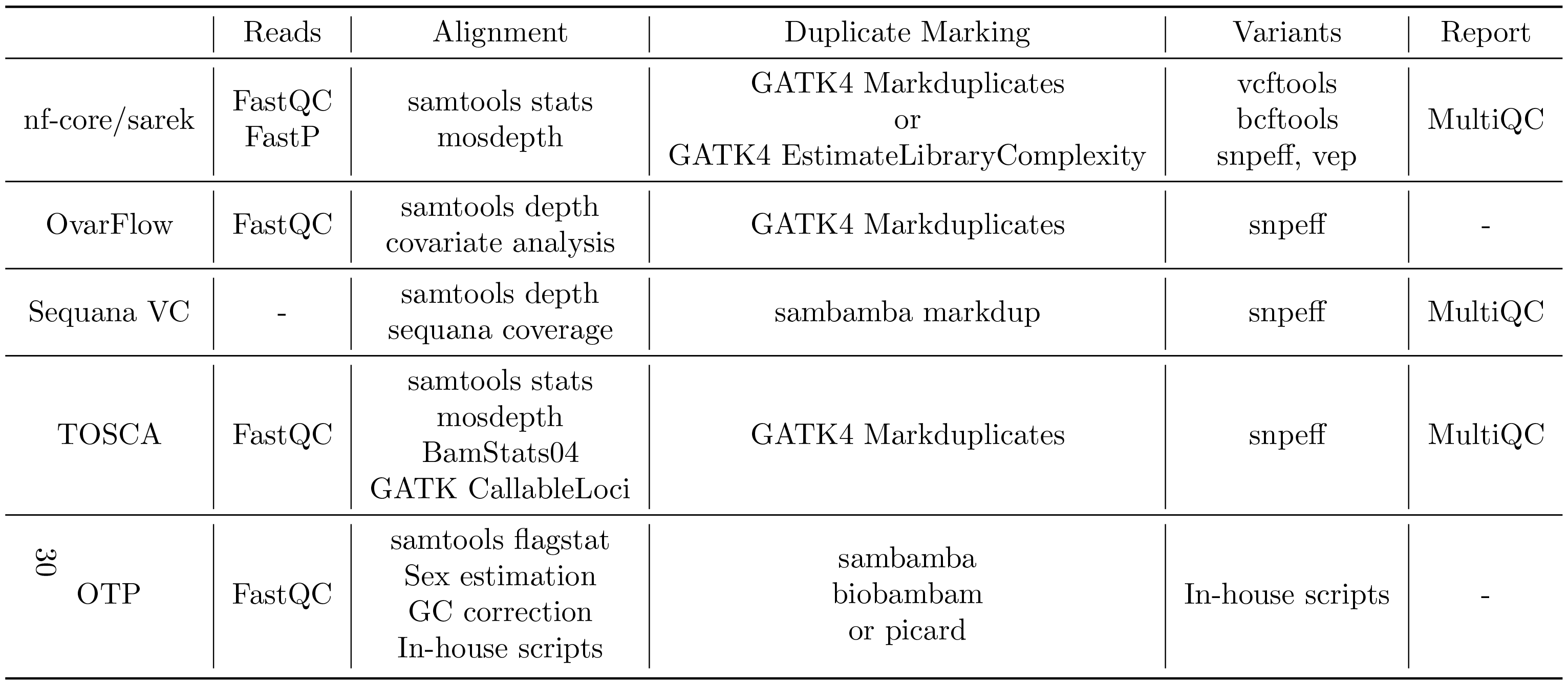
Variant calling pipelines typically use a set of quality control tools to allow introspection of data quality and intermediate analysis steps. Here, the various tools used by the pipelines described in table 1 are shown. All pipelines report basic mapping, duplication metrices, and variant calling metrics. nf-core/sarek, sequana VC, and TOSCA combine these into a MultiQC report.

## Appendix B Intra-sample parallelization

**Fig. B1:**
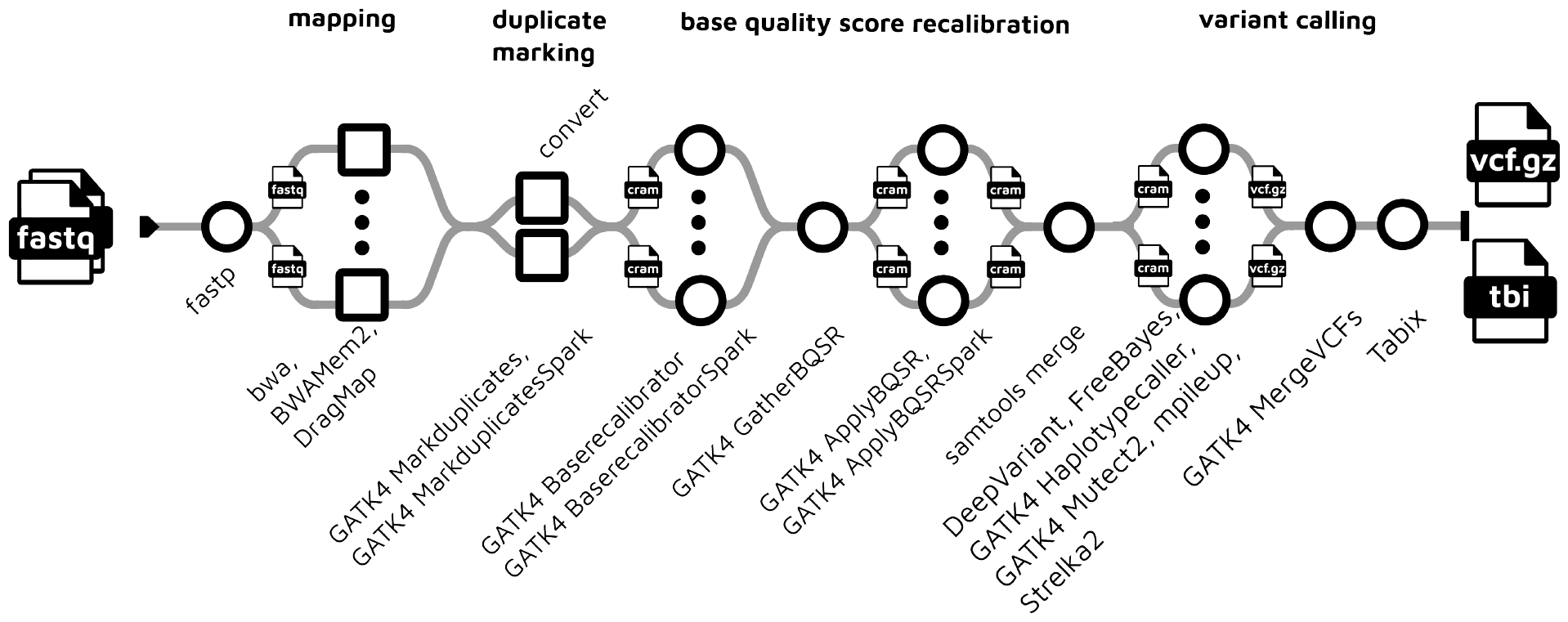
The figure depicts the processing of a single sample. While Nextflow runs the analysis of each sample in parallel, intra-sample parallelization was implemented for the mapping step by splitting the FastQ files beforehand, mapping each, and merging all FastQ files belonging to one sample and duplicate marking them. Base quality score recalibration and SNP/Indel calling are run across genomic intervals, which can be either user-provided or GATK-provided interval files.

## Appendix C Table with detailed changes between Sarek 2.5.2 and 3.1.1

**Table C2:**
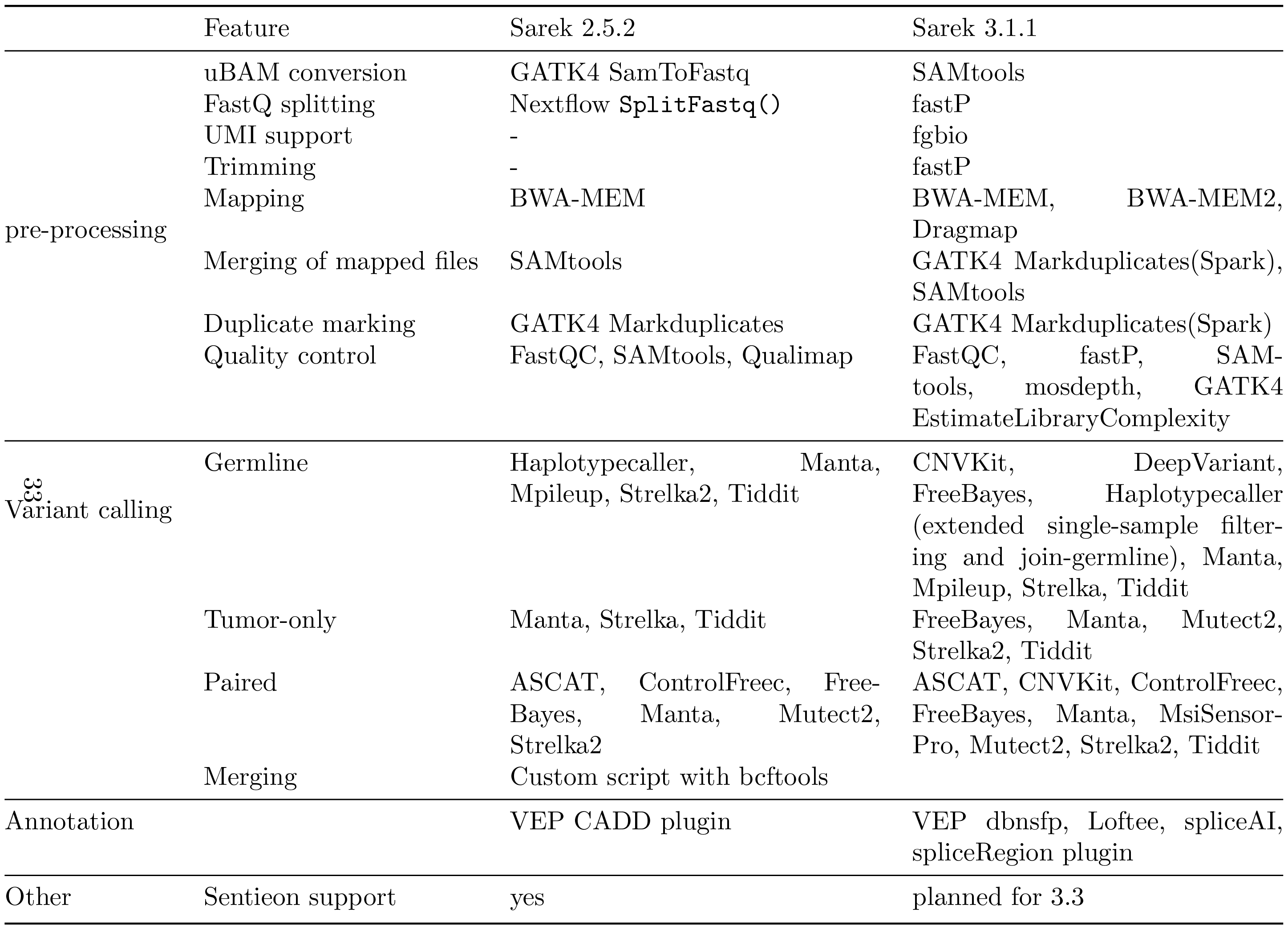
The updates since nf-core/sarek 2.5.2 are summarized in this table: most notably adapter trimming and UMI support were added, and read splitting was optimized and enabled for large FastQ files by using fastP. Merging of aligned BAM files across splits and lanes was incorporated into the duplicate marking process. Furthermore, individual steps allow a wider tool selection, i.e. BWA-MEM2 and Dragmap for mapping and further tools for variant calling. Sentieon support for all steps will be added with release 3.3.

## Appendix D Reducing storage requirements with CRAM

**Fig. D2:**
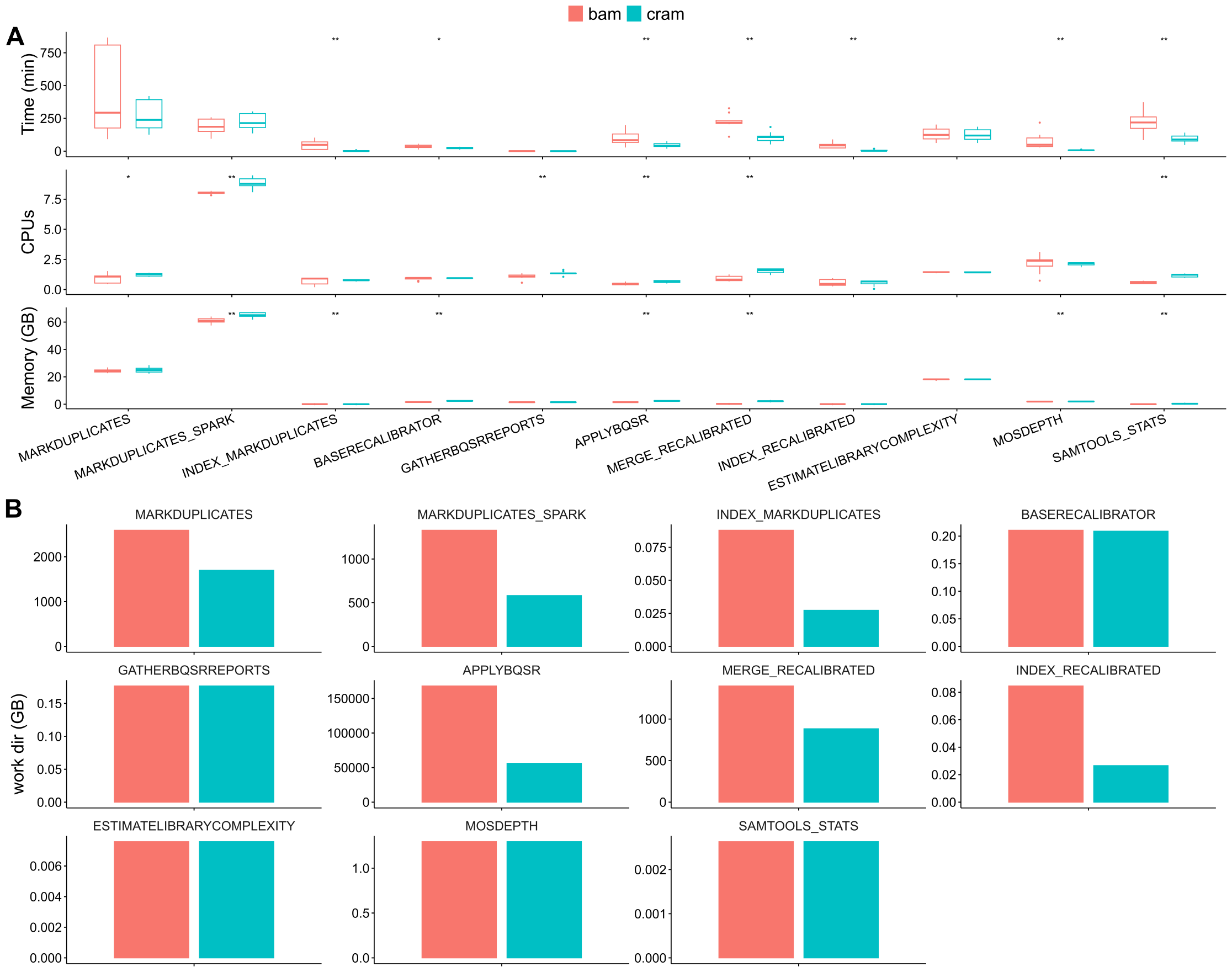
Resource usage of nf-core/sarek 3.1.1 when storing intermediate data in BAM versus CRAM format for all pre-processing processes. **A)**: Average realtime, and maximum CPU and memory usage (peak rss) as reported by the Nextflow trace file. For processes split within a sample (i.e. ApplyBQSR), the task with the highest runtime per sample is shown as the process runtime. Resource usage was compared using the paired Wilcoxon test (^**^ *p <* 0.01, ^*^ *p <* 0.05). Seven of the eleven processes are significantly faster when using the CRAM format. Five processes have a significantly higher CPU hour usage, and three require more memory in comparison to using BAMs. **B)**: Storage was evaluated by calculating the total size of the work directories of all tasks of the respective process. The storage usage between both is identical, due to Nextflow accessing the input files via symlinks. Thus only the output is measured here for each process, which is independent of input format. Each condition was repeated three times for samples of five tumor-normal paired patients.

**Fig. D3:**
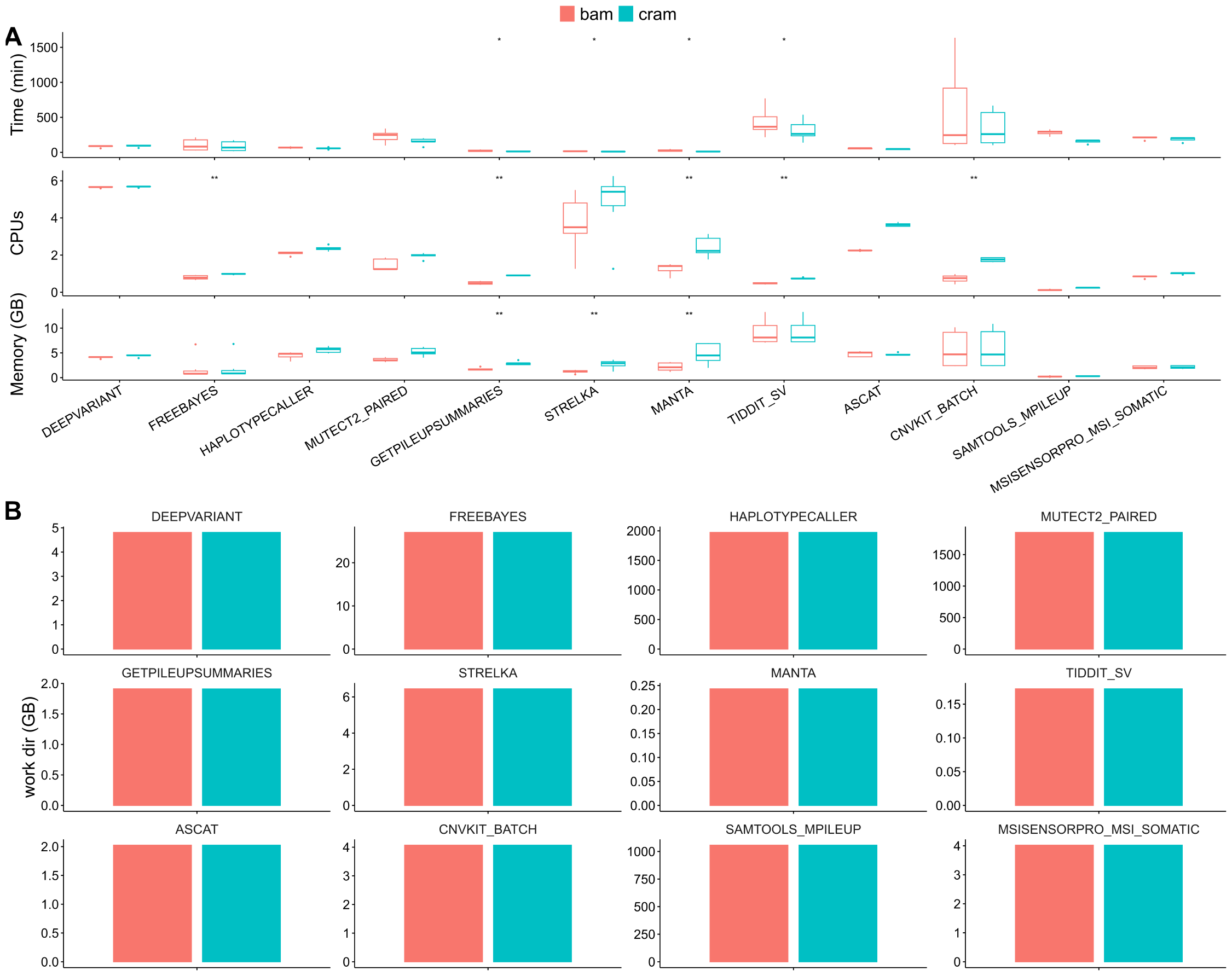
Resource usage of all variant calling processes in nf-core/sarek 3.1.1 when starting from BAM and CRAM format. **A)**: Average realtime, and maximum CPU and memory usage (peak rss) as reported by the Nextflow trace file. For processes split within a sample, the task with the highest runtime per sample is shown as the process runtime. Resource usage was compared using the paired Wilcoxon test (^**^ *p <* 0.01, ^*^ *p <* 0.05). Four of the 12 processes are significantly faster when using the CRAM format. Six have a significantly higher CPU hour usage, and seven require more memory in comparison to using BAMs. **B)**: Storage was evaluated by calculating the total size of the work directories of all tasks of the respective process. Six processes have reduced storage requirements. Each condition was repeated three times for samples of five tumor-normal paired patients.

## Appendix E Sharding FastQ files reduces runtime

**Fig. E4:**
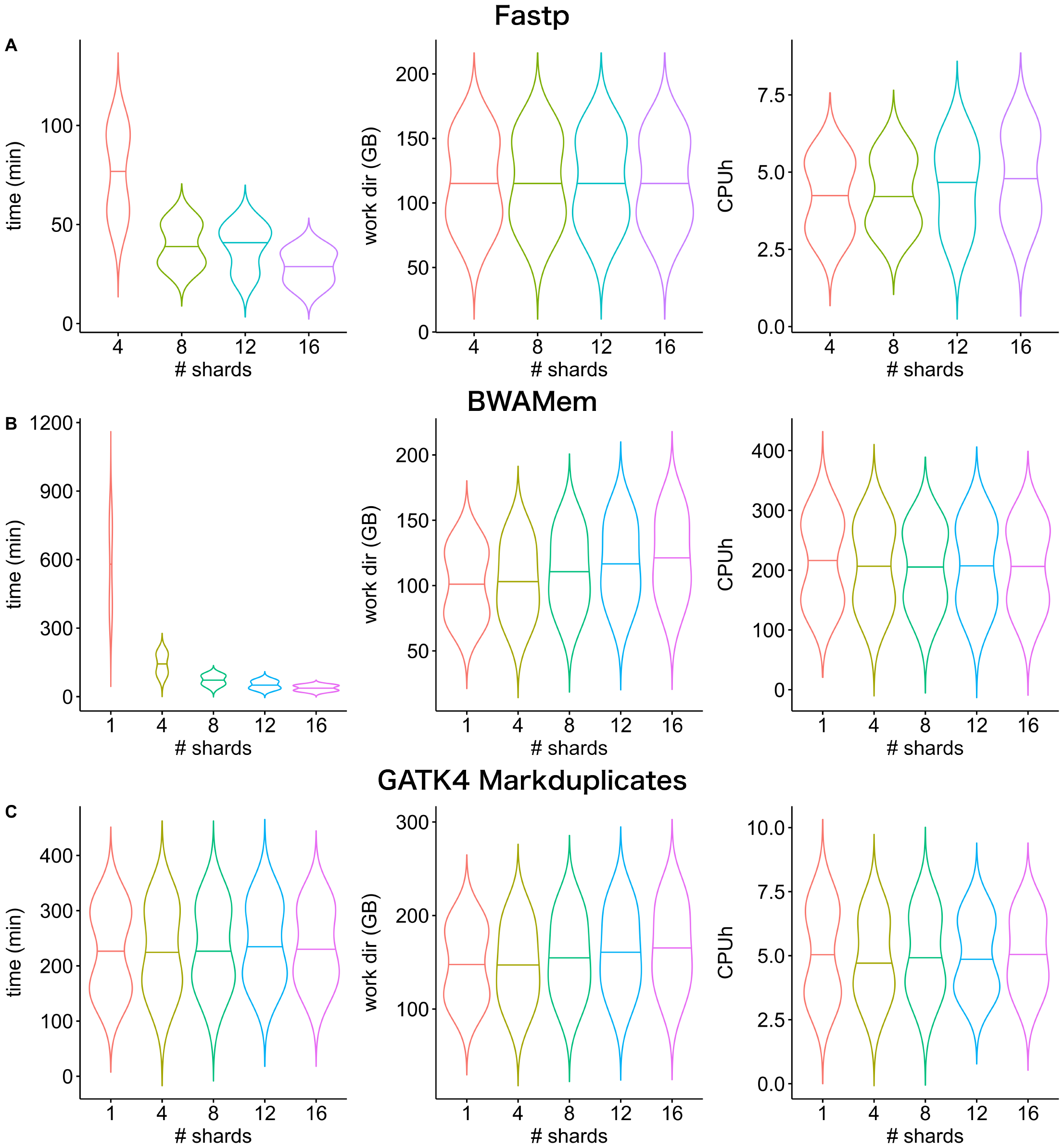
Dividing the input FastQ files into increasing amounts of shards. **A)**: Resource usage of fastP during sharding of the input FastQ files. The tool was run with a different count of CPUs corresponding to the desired number of shards. **B)**: Resource usage of BWA-MEM during mapping of each shard. **C)**: Resource usage of the duplicate marking process. Merging of sharded bam files and duplicate marking is performed with GATK4 Markduplicates. CRAM conversion is done with SAMTools. The violin plots show computations on tumor-normal paired samples of five patients. The time was evaluated by summing up the highest realtime per task per sample as reported by the Nextflow trace report. The work directory size and CPU hours are the sums of all involved tasks.

## Appendix F Splitting by intervals reduces runtime

**Fig. F5:**
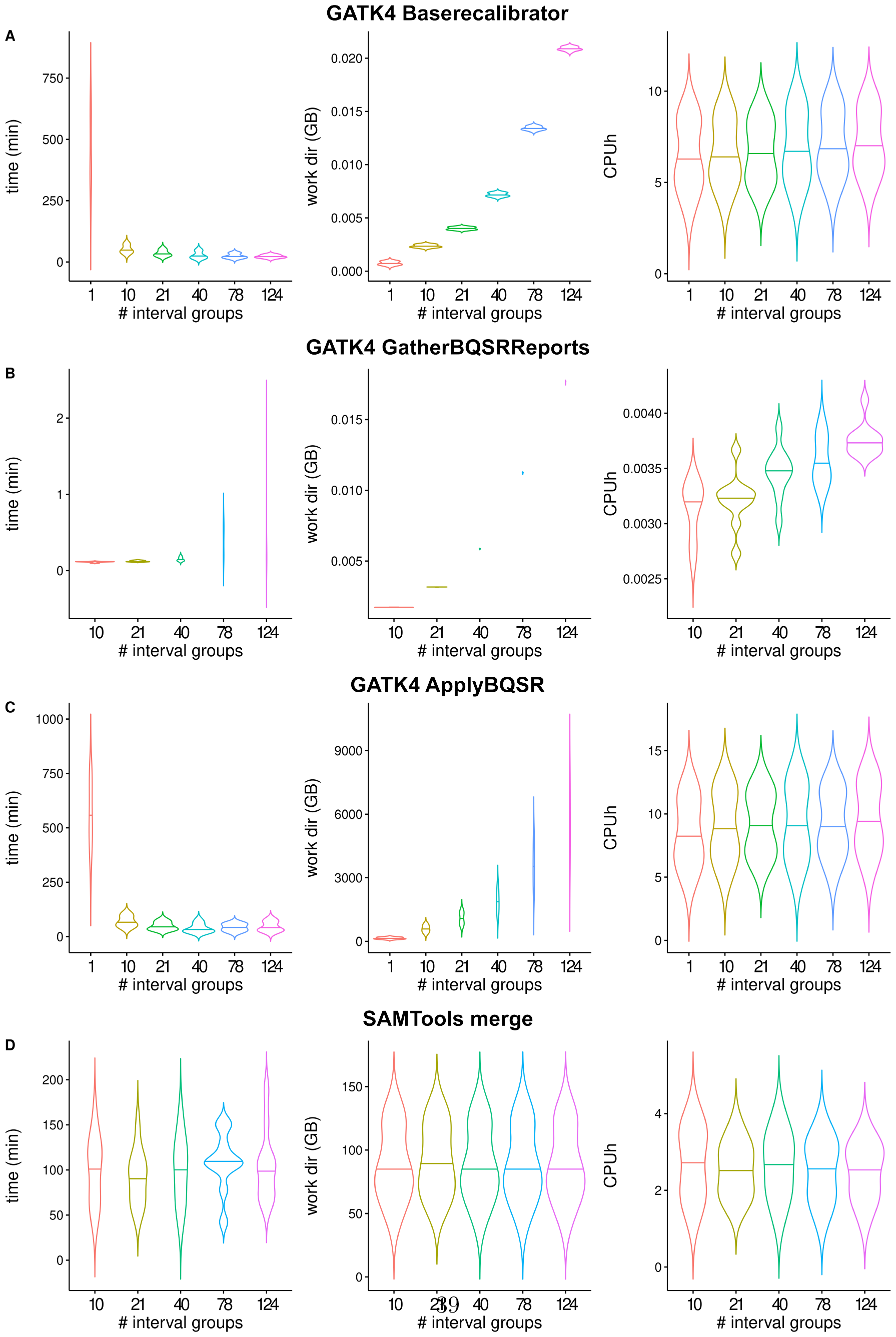
Parallel processing of interval groups reduces runtime **A)**: GATK4 BaseRe-calibrator runtime decreases with an increasing number of interval groups. Storage space requirements increase, while CPU hours stay consistent. **B)**: For GATK4 Gather-BQSRReports all three metrics increase with an increasing number of interval groups. **C)**: For GATK4 ApplyBQSR the runtime decreases with an increasing number of interval groups. Storage space requirements increase, while CPU hours stay consistent. **D)**: For SAMTools Merge all three metrics are consistent across the number of interval groups. The violin plots show computations on tumor-normal paired samples of five patients for each tool. The time was evaluated by summing up the highest realtime per task per sample as reported by the Nextflow trace report. The work directory size and CPU hours are the sums of all involved tasks.

**Fig. F6:**
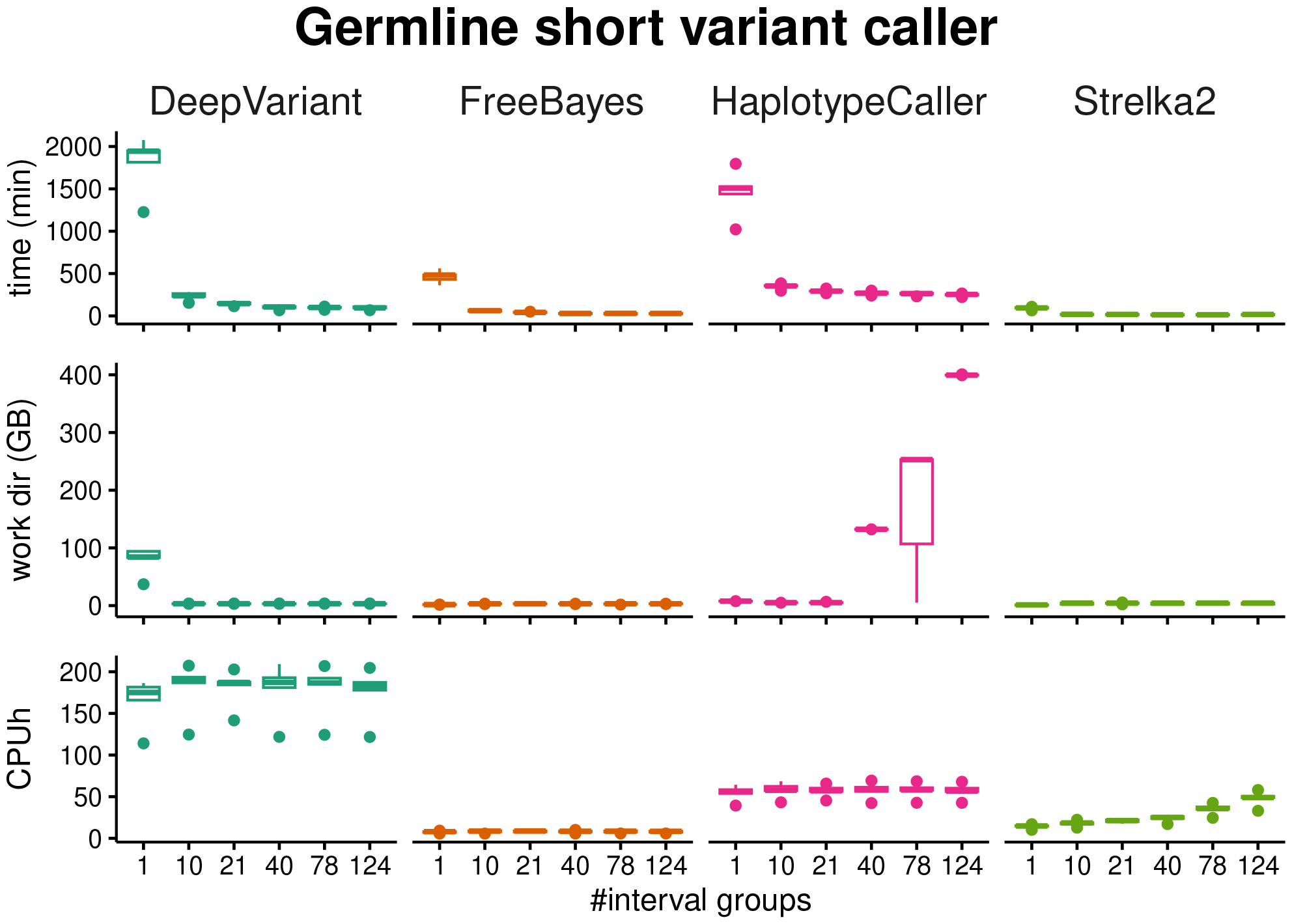
Effect of parallelizing computations across interval groups on germline variant calling processes, which include the respective variant caller followed by GATK4 MergeVCFs. FreeBayes VCFs are sorted before merging. GATK4 HaplotypeCaller is followed by GATK4 CNNSCoreVariants and GATK4 FilterVariantTranches. All variant callers speed up on parallel processing across 10 interval groups. DeepVariant and HaplotypeCaller speed up further with fewer interval groups. Storage usage decreases for DeepVariant for 10 interval groups and remains stable. FreeBayes and Strelka2 have similar storage usage across parallelization. For VCFs called by HaplotypeCaller storage usage increases. CPU hours are similar across degrees of parallelization with an increase measured for Strelka2. The violin plots show computations on normal samples of five patients. The time was evaluated by summing up the highest realtime per task per sample as reported by the Nextflow trace report. The work directory size and CPU hours are the sums of all involved tasks.

**Fig. F7:**
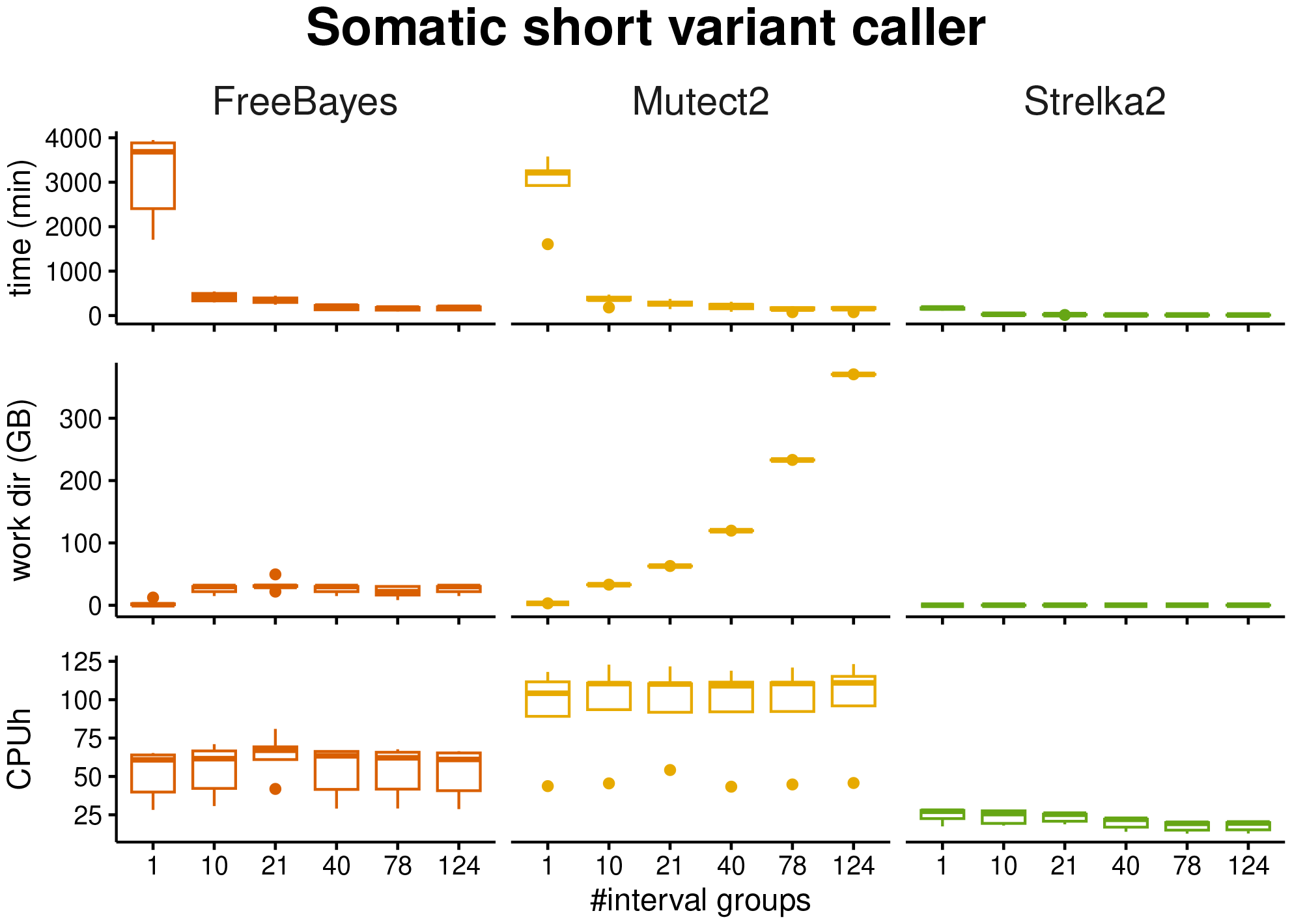
Effect of parallelizing computations across interval groups on somatic variant calling processes, which include the respective variant caller followed by GATK4 MergeVCFs. FreeBayes VCFs are sorted before merging. All variant callers speed up on parallel processing across 10 interval groups. FreeBayes and Mutect2 further speed up with 21 interval groups. Storage usage increases for FreeBayes for 10 interval groups and remains stable. For Mutect2 storage usage increases with increasing interval groups. It remains stable for Strelka2. CPU hours are similar across degrees of parallelization with a decrease measured for Strelka2. The violin plots show computations on tumor-normal paired samples of five patients. The time was evaluated by summing up the highest realtime per task per sample as reported by the Nextflow trace report. The work directory size and CPU hours are the sums of all involved tasks.

## Appendix G Benchmarking against truth datasets

**Fig. G8:**
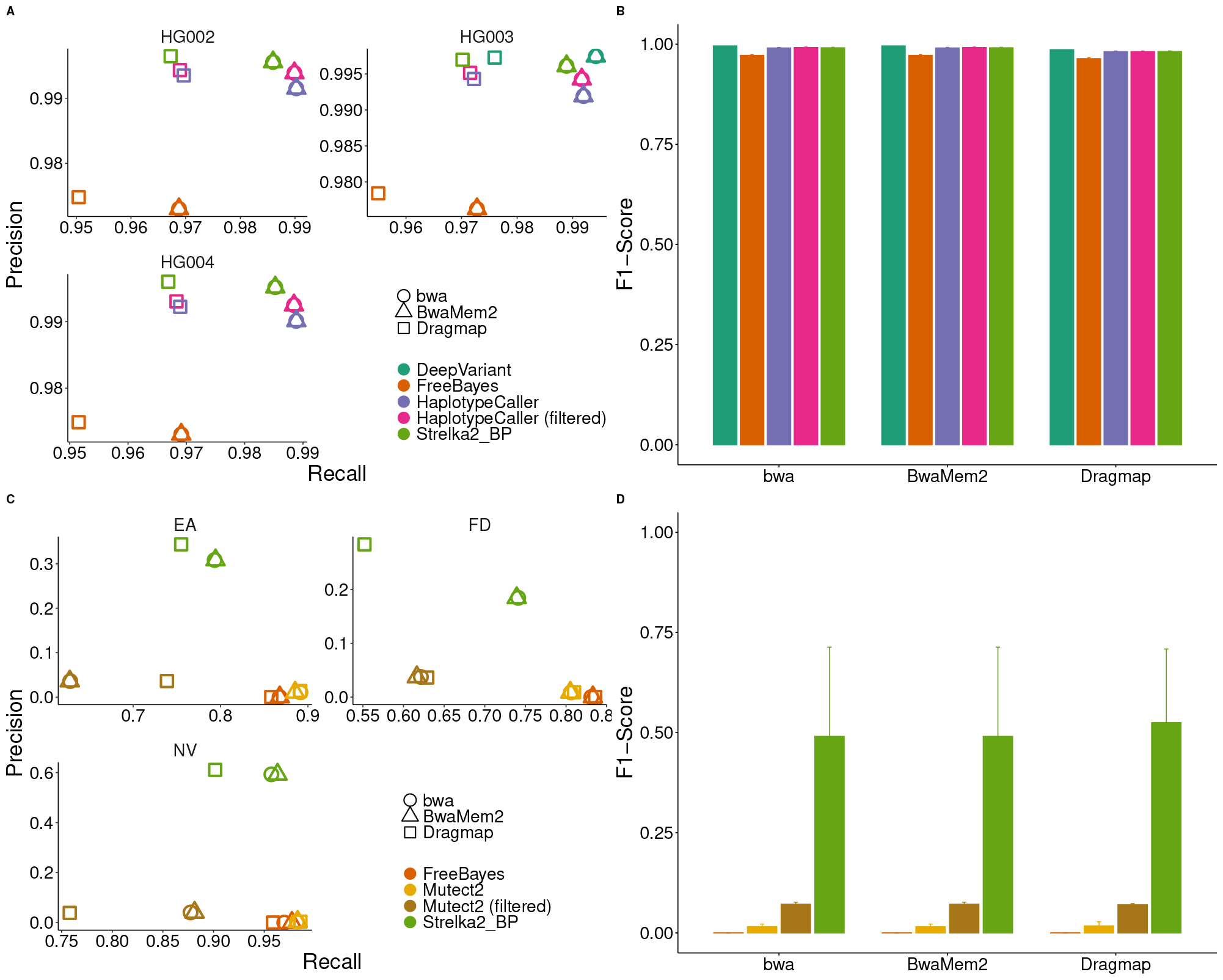
Germline and somatic variant calling evaluation of high-confidence calls using ground-truth benchmarking data with respect to Indels. **A**,**B**: The germline track of the pipeline was evaluated using 3 WGS GiaB datasets (HG002-HG004). The average precision, recall and F1-score values across all the samples are plotted respectively. **C**,**D**: The paired calling track was evaluated using three tumor-normal WES pairs (EA, FD, NV) from SEQ2C.

**Fig. G9:**
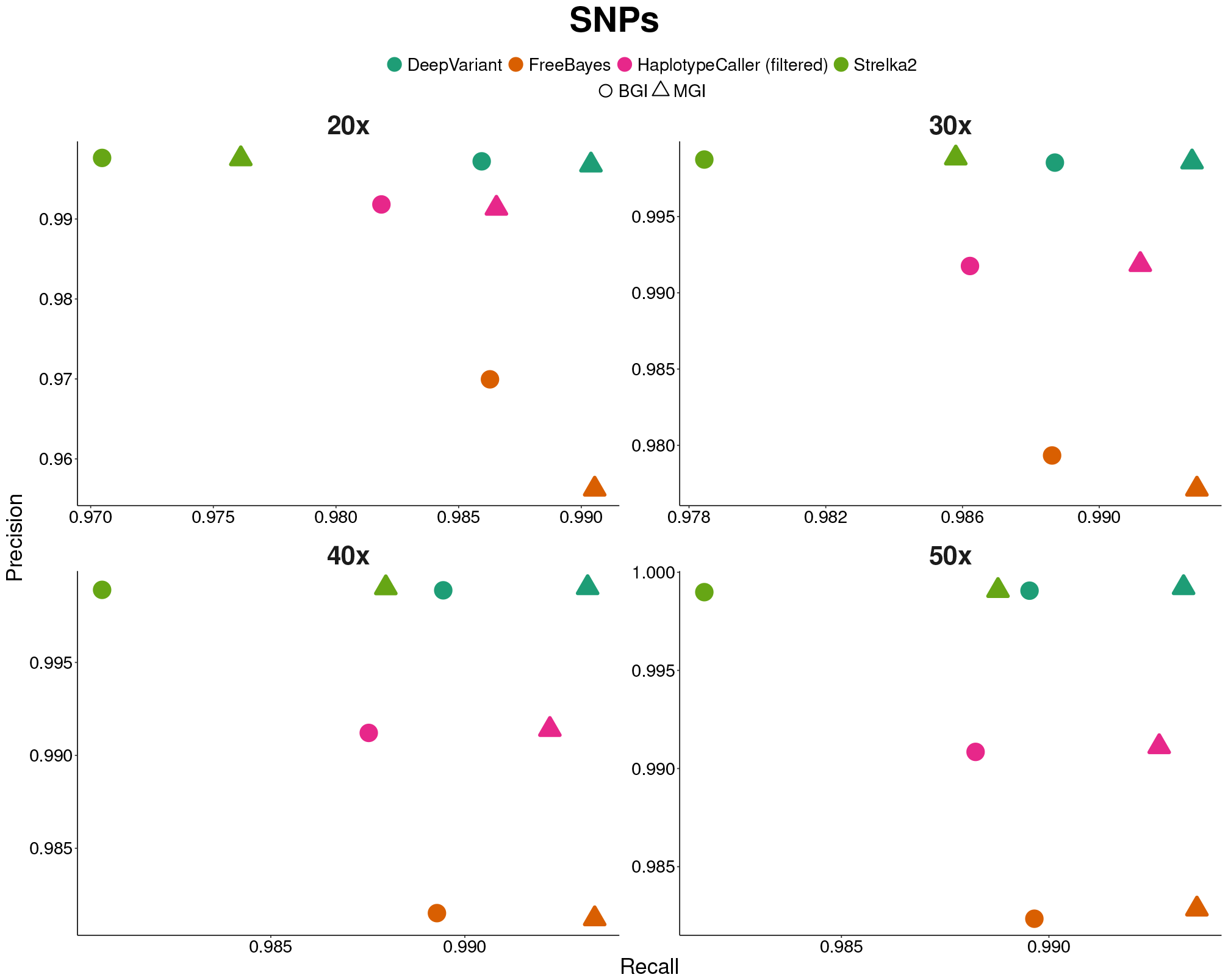
Germline variant calling evaluation of high-confidence calls using ground-truth benchmarking data with respect to SNPs. Samples from MGISeq and BGISeq500 were mapped with BWA-MEM. Different coverages were used as input. For all investigated coverage values FreeBayes and DeepVariant have the highest recall. Strelka2 and DeepVariant show the highest precision values for all samples.

**Fig. G10:**
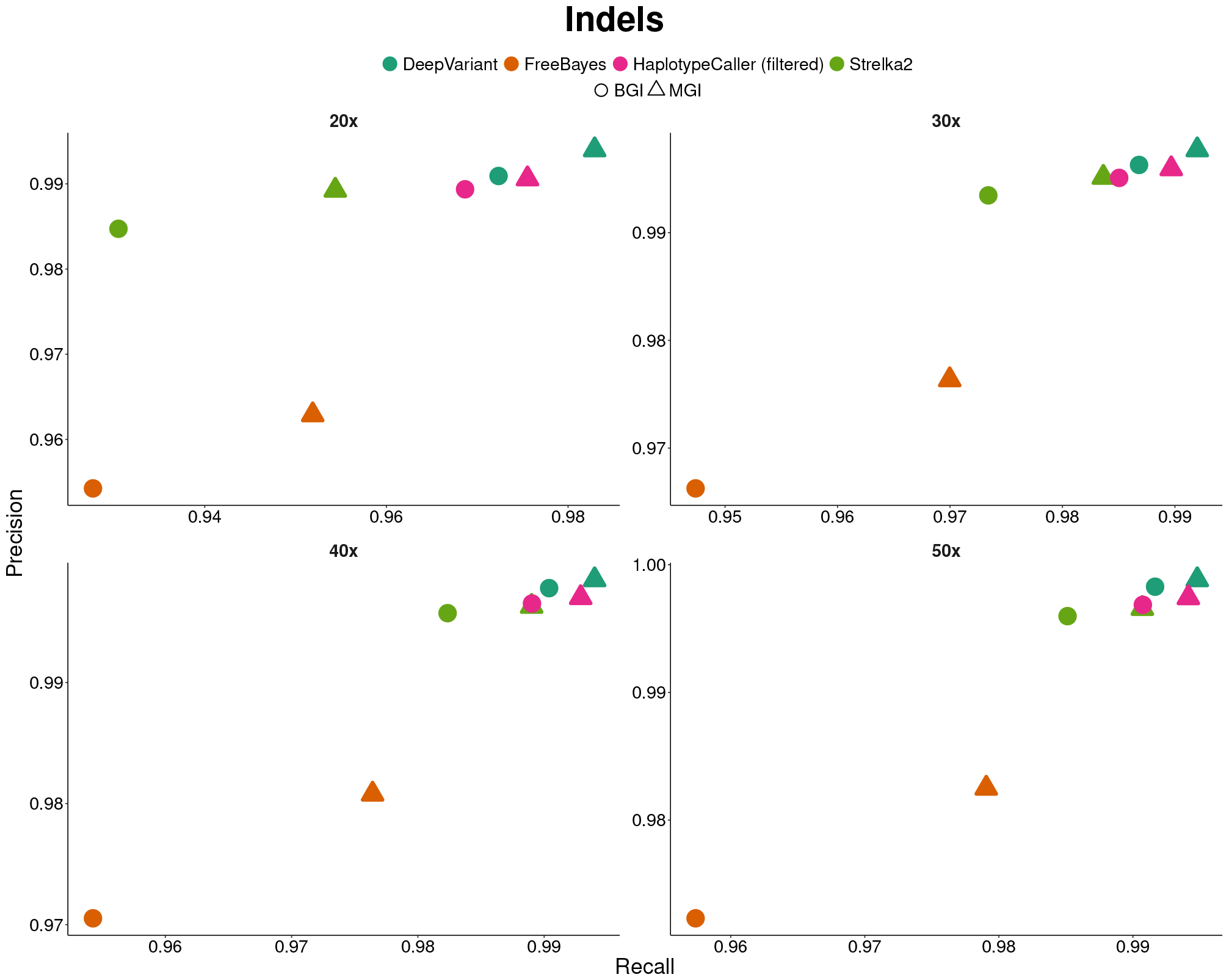
Germline variant calling evaluation of high-confidence calls using ground-truth benchmarking data with respect to Indels. Samples from MGISeq and BGISeq500 were mapped with BWA-MEM. Different coverages were used as input. For all investigated coverage values HaplotypeCaller and DeepVariant have the highest recall, followed by Strelka2. MGI samples analyzed with DeepVariant had the highest precision values.

## Appendix H Comparison of CNV calls against PCAWG samples

**Fig. H11:**
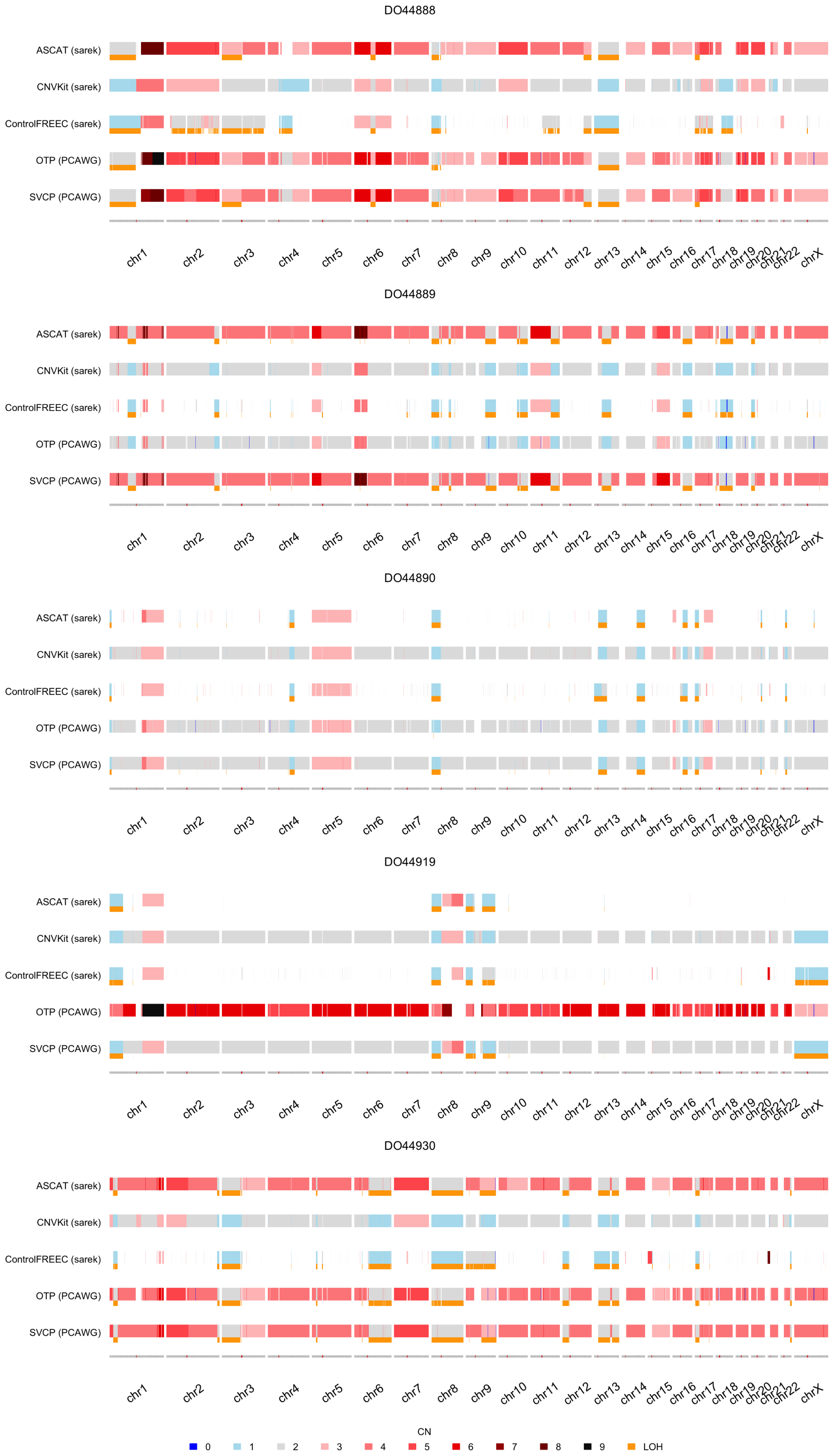
Comparison of copy number calls obtained with nf-core/sarek using ASCAT, Control-FREEC, and CNVKit to the ones from the PCAWG study downloaded from the ICGC portal. For the latter, there are two results files available for each patient respectively, one called with the OTP pipeline, one called with the Sanger pipeline. The PCAWG calls agree for 3 patients. For DO44890 all calls across all pipelines and tools agree. For DO44930 and DO44888, the nf-core/sarek ASCAT calls are similar to both the OTP and Sanger pipeline based calls, the CNVKit and Control-FREEC calls differ, but are similar towards each other. For DO44889 the nf-core/sarek and Sanger pipeline calls overlap, as well as the OTP calls and nf-core/sarek Control-FREEC and CNVKit calls. Lastly, for DO44919, all nf-core/sarek calls overlap with the Sanger pipeline results, the OTP pipeline results differ.

## Footnotes

1 https://github.com/fulcrumgenomics/fgbio

2 https://github.com/Illumina/DRAGMAP)

3 https://gatk.broadinstitute.org/hc/en-us/articles/360035889551-When-should-I-restrict-my-analysis-to-specific-intervals-

4 https://github.com/Illumina/strelka/blob/v2.9.x/docs/userGuide/README.md

5 https://github.com/Ensembl/VEP_plugins/blob/release/109/SpliceRegion.pm

6 https://github.com/FriederikeHanssen/sarek/tree/bam_31

7 https://seqera.io/blog/the-state-of-the-workflow-2023-community-survey-results/

8 https://seqera.io/blog/breakthrough-performance-and-cost-efficiency-with-the-new-fusion-file-system/

9 https://github.com/nf-core/configs/blob/c709be3b599d463fcfa82196fd4c9c5fa1e99513/conf/cfc.config

10 https://github.com/lh3/seqtk

11 https://github.com/illumina/hap.py

12 https://cloud.tower.nf/

